# LEADER (Leaf Element Accumulation from Deep Roots): a nondestructive phenotyping platform to estimate rooting depth in the field

**DOI:** 10.1101/2023.05.02.539074

**Authors:** Meredith T. Hanlon, Kathleen M. Brown, Jonathan P. Lynch

## Abstract

Deeper rooted crops are an avenue to increase plant water and nitrogen uptake under limiting conditions and increase long-term soil carbon storage. Measuring rooting depth, however, is challenging due to the destructive, laborious, or imprecise methods that are currently available. Here, we present LEADER (Leaf Element Accumulation from DEep Roots) as a method to estimate in-field root depth of maize plants. We use both X-Ray fluorescence spectroscopy (XRF) and ICP-OES (Inductively Coupled Plasma Optical Emission spectroscopy) to measure leaf elemental content and relate this to metrics of root depth. Principal components of leaf elemental content correlate with measures of root length in four genotypes (R^2^= 0.8 for total root length), and we use linear discriminant analysis to classify plants as having different metrics related to root depth across four field sites in the United States. We can correctly classify the plots with the longest root length at depth with high accuracy (accuracy greater than 0.6) at two of our field sites (Hancock, WI and Rock Spring, PA). We also use strontium (Sr) as a tracer element in both greenhouse and field studies, showing that elemental accumulation of Sr in leaf tissue can be measured with XRF and can estimate root depth. We propose the adoption of LEADER as a tool for measuring root depth in different plant species and soils. LEADER is faster and easier than any other methods that currently exist and could allow for extensive study and understanding of deep rooting.

## 1. Introduction

Rooting depth has important effects on the acquisition of soil resources, interactions with soil biota, plant anchorage, and atmospheric carbon sequestration (Lynch and Wojciechowski 2015; Thorup-Kristensen et al. 2020). Rooting depth is also exceedingly difficult to measure in soil. Rooting depth can be easily measured in (typically small) plants grown in artificial media, the most extreme example being humid air (‘aeroponics’ (Eldridge et al. 2020; Awika et al. 2021)), but also commonly including agar plates (Guseman et al. 2016; Ogura et al. 2019), solution culture (Mathieu et al. 2015), paper growth pouches (Atkinson et al. 2015), or systems in which roots grow in solid media against a transparent surface (Nagel et al. 2012; Rellán-Álvarez et al. 2015; Yuan et al. 2016; Deja-Muylle et al. 2021; LaRue et al. 2022). Such systems are useful in specific contexts but may not be correlated with root phenotypes observed in natural soil, which has many characteristics capable of influencing root growth including, but not limited to, mechanical impedance, soil biota, and hypoxia.

The measurement of root depth in soil faces several challenges, including 1) root systems are highly complex objects composed of many distinct elements, many of which are small; 2) the distribution of roots with soil depth is dynamic in space and time; 3) spatiotemporal variability in soil properties within and among soil taxa can generate substantial variation in root phenotypes because of root plasticity; and 4) soil is a heterogenous, dense, and opaque medium. Direct observation of rooting depth in soil is, therefore, challenging. No methods exist that combine high throughput, accuracy, use readily accessible technology, and are not destructive of the plant or the research plot (Taylor et al. 2013; Maeght et al. 2013; Burridge et al. 2020; Cabal et al. 2021). While a comprehensive review of methods for root measurement in the field is beyond the scope of this article, a very brief summary of the merits and limitations of common methods follows.

### Direct excavation

Monolith excavation of entire root systems is possible (Böhm and Köpke 1977; Wu and Guo 2014; Ning et al. 2015; Teramoto et al. 2019) but only with substantial effort, likely displacement of the original position of axial roots, loss of small diameter roots in soil washing, and substantial destruction of the plant as well as substantial disturbance of the field plot. Trench excavation exposes a soil surface that represents a 2D transect of root distribution (Nielsen et al. 1999; van Noordwijk et al. 2000) but is also laborious and destructive. Excavation of the root crown in *Shovelomics* methods does not provide measurement of root depth *per se* (Trachsel et al. 2011; Colombi et al. 2015).

### Soil coring

Soil coring enables a direct measurement of root distribution with depth, is moderately laborious, requires simple equipment, and does not require destruction of the subject plant or substantial damage to a research plot. However, because of the relatively small size of soil cores and the spatiotemporal complexity of root distribution, a large number of samples are required to estimate rooting depth in a field plot, especially in row crops such as maize or grain legumes with substantial interrow spacing (Taylor et al. 2013; Burridge et al. 2020).

### Minirhizotrons

Transparent minirhizotron observation tubes permit nondestructive observation of roots encountering the outside of the tube as a function of depth and time (Taylor 1987). Once installed they are relatively easy to use, but the large number of images generated require significant analytical resources, and more importantly, the interface of the tube and the soil may not be representative of root distribution in bulk soil, as is true of any measurement system that requires roots to grow against a transparent surface. This is especially a concern in the context of soil mechanical impedance, which can be quite different from bulk soil as a result of tube placement and/or soil shrinkage or expansion in response to changes in soil water content. Centralized facilities constructed with many minirhizotron observation tubes placed throughout the profile are useful (Svane et al. 2019; Rajurkar et al. 2022) but only sample the soil and environment at one location.

### Non-invasive methods

Magnetic resonance imaging (MRI), X-ray computed tomography (CT), and positron emission tomography (PET) permit non-destructive measurement of root distribution (Schulz et al. 2012; Mooney et al. 2012; Mairhofer et al. 2013; Pfeifer et al. 2015; Wasson et al. 2020) but suffer tradeoffs between sample volume and spatial resolution, and require complex, bulky, and costly instrumentation, so as yet are not suited to field work. Ground penetrating radar (GPR) is being used for coarse roots of trees (Hruska et al. 1999; Alani and Lantini 2020) and has agronomic applications to the coarser roots of root and tuber crops (Guo et al. 2013; Wu et al. 2014; Delgado et al. 2017) but is not yet capable of detecting fine roots, which are responsible for the majority of water and nutrient acqusition. New indirect methods include electrical-impedance spectroscopy (Peruzzo et al. 2021), in-field MRI (Bagnall et al. 2020), thermoacoustic imaging (Singhvi et al. 2022), ultra-wideband microwave imaging (Shi et al. 2022), electrical resistance tomography (Srayeddin and Doussan 2009; Whalley et al. 2017), though all of these technologies are emerging. They are also depth-limited based on the probe size and may not work in all soils.

### Tracer application

Indirect measurement of rooting depth is possible by observing shoot responses to tracers placed at specific soil depths. For example, differential injury from herbicides placed at a specific soil depth has been used to distinguish deeper rooted crop genotypes in the greenhouse (Al-Shugeairy et al. 2014) and field (Corre-Hellou and Crozat 2005). Detection of naturally occurring or added tracers such as oxygen isotopes, heavy water or deuterium offer estimates of depth of water acquisition (Chimungu et al. 2014a, b; Kulmatiski et al. 2017; Rothfuss and Javaux 2017; Chen et al. 2019, 2021). Shoot accumulation of ^15^N from pulses injected at specific soil depths has been used to estimate genotypic variation in the depth of N acquisition in maize (Saengwilai et al. 2014b; Chimungu et al. 2015; Chen et al. 2019; Schneider et al. 2022; Wacker et al. 2022). Other tracers, including Cs, Li, Rb, Sr, and Se have been used and applied through in-growth cores, which alter the soil environment and may impact root growth (Rasmussen et al. 2020). These methods are useful but are limited by the transient nature of the injected tracer, and the difficulty of using them to compare many plots, as is needed for example in breeding or genetic studies.

In this article we present LEADER (Leaf Element Accumulation from DEep Roots), a nondestructive method to estimate rooting depth in the field through measurement of foliar elemental accumulation. The bioavailability of soil minerals, including mineral nutrients as well as non-nutrient elements, varies with soil depth in natural soils. In most soils, soil morphogenesis results in horizons with distinct physical, chemical, and biological characteristics. In young soils with minimal horizonation, variation of temperature, water content, oxygen availability, organic matter deposition, and biotic activity with depth create differences in the bioavailability of soil elements for root uptake. As roots explore the soil, they actively acquire mineral nutrients and passively accumulate any soluble elements at the root tip, which lacks apoplastic barriers. Elements passively entering the root tip through the apoplast may eventually be transported to the shoot through transpiration. Elemental accumulation in shoot tissue therefore represents a bioassay of elemental bioavailability in the soil, which should therefore be correlated with rooting depth. In this article we present a proof of concept of LEADER by showing correlation of rooting depth assayed through soil coring to variation in rooting depth in maize genotypes grown in several natural soils using handheld X-Ray Fluorescence (XRF) Spectroscopy to measure foliar elemental content. We find that LEADER is effective in differentiating between deeper and shallower rooting maize lines in the field.

## 2. Materials and Methods

### 2.1 Field LEADER experiments

Field experiments were conducted in the summer of 2020 at the Russell E. Larson Agricultural Research Center in Rock Spring, PA, USA (40.710272, -77.952788). All seeds were provided by Shawn Kaeppler at the University of Wisconsin, Madison. Four genotypes (inbred lines B73 and PHB47 and hybrid lines DKC49 x 94RIB and DKC35 x 88RIB) were planted in 16 rows 65 m in length with 76 cm row spacing and 16.5 cm between plants (79,00 plants ha^-1^). Supplemental nitrogen was provided in the form of urea at 200 kg ha^-1^. No supplemental irrigation was supplied. At six, eight, and ten weeks after germination, plants and soil were sampled. Four plants of each genotype were selected based on having a representative growth habit, low weed pressure, and sufficient germination of neighboring plants on all sides. All leaves were harvested and oven-dried for later analysis. Cores were obtained for soil nutrient analysis and root depth distribution by driving a steel coring tube of 5 cm diameter to a depth of 60 cm using sledgehammers. Root distribution cores were taken 5 cm from the base of the plant of interest. Soil cores were taken between rows near the plants of interest. Cores were stored in plastic tubing under refrigeration at 4°C before being separated into 10 cm segments. For root length distribution, roots were washed from the cores using low pressure water over a 2 mm sieve. Roots were collected into 70% ethanol and stored at 4°C. Roots were after scanned on an Epson Perfection V850 (Epson America) scanner at 300 DPI with the roots held in a plastic tray with water and analyzed via WinRHIZO (Regent Instruments). Soil samples were air-dried, placed in plastic bottles, and shaken on a paint shaker (Sport DC-1-C, Miracle Paint, Inner Grove Heights, MN) until homogenized. Samples were sieved through a 2 mm sieve and subsequently analyzed for elemental content.

### 2.2 Field extrapolation experiments

In the summer of 2019, a set of 30 maize genotypes were grown at four locations across the US. These locations were the Russell E. Larson Agricultural Research Center in Rock Spring, PA, USA (40.710272, -77.952788), the University of Colorado Agricultural Research and Education center (40.6502515, -104.9953871) in Fort Collins, CO, and two field research stations in Wisconsin: The University of Wisconsin Arlington Agricultural Research Station (43.303344, - 89.386564) and the University of Wisconsin Hancock Agricultural Research Station (44.118611, -89.555278). In 2020, 50 hybrid maize genotypes were planted at the Russell E. Larson Agricultural Research Center in Rock Spring, PA. We selected 9 of these lines for analysis. Stress treatments were imposed in PA, CO, and Hancock WI. Plants in PA were grown under low nitrogen conditions; CO, under water-deficit conditions; and Hancock, low nitrogen, water deficit, and the combination of the treatments. Full information for these locations, including field management and soil characteristics can be found in Supplemental Table 1. Information on the germplasm used at each site can be found in Supplemental Table 3.

We selected between 6 and 12 genotypes to collect root depth information via coring using a steel coring tube of 5 cm diameter to a depth of 60 cm. The tube was driven into the ground either with sledgehammers or a tractor-mounted hydraulic Giddings rig (Giddings Machine Company, Windsor, CO). Cores were obtained after flowering in all locations. Three cores were taken in-row from each plot and were stored at 4°C until they were processed for root length information using the methods above. Ear leaves were collected and bagged from one plant per plot taken near the site of coring. Ear leaves were dried at 60°C before being analyzed for elemental content via XRF.

### 2.3 Greenhouse experiments

Greenhouse experiments were conducted in 2018 and 2019 to test the efficacy of strontium as a tracer for LEADER analysis. All seeds were provided by Shawn Kaeppler at the University of Wisconsin Madison. Greenhouse pot experiments were conducted in a greenhouse at University Park, PA (40.802336, -77.862308) Day length was 14 h, and the temperature was maintained at approximately 28°C/26°C day/night and natural light levels were supplemented with LED light when photosynthetic active radiation dropped below 900 m^−2^ s^−1^. All plants were grown in PVC mesocosms (15.5 cm diameter, 1.6 m height) lined with a 6 mm high-density polyethylene film. The 30L mesocosms were filled with 50% sand (medium grain, Q-ROK, US Silica), 30% vermiculite (D3 fine, Whittemore), 5% perlite (Supercoarse 23, Whittemore) and 20% soil. Soil was collected from the top 20 cm of soil at the Russell E. Larson Agricultural Research Center at Rock Spring, PA, air-dried, crushed, and sieved through a 4-mm mesh.

In 2018, a single inbred maize genotype, Mo17, was used. The media mix included 70g of Osmocote Plus slow-release fertilizer (15-9-12). Mesocosms were filled with 11.25 L of media until media reached 100 cm from the top and the media was saturated with water. We then saturated 5 L of media with 600 mL of a treatment resulting in a treatment layer concentration of 0.44 mM SrCl_2_, 1.3 mM SrCl_2_, 2.6 mM SrCl_2_, 0.44 mM RbCl, or 1.3 mM Rb and water for the control treatments. The mesocosms were then filled to the top, saturated with water, two seeds were planted per mesocosm, and they were thinned to one plant after germination. Four columns were randomized into each treatment, and columns were irrigated with 200 mL of water daily for two weeks and twice daily afterwards. Plants were scanned using handheld XRF (see below, 2.5 XRF Analysis) at 28, 34, and 42 days after planting. At 43 days, plants were harvested, and shoot tissue was dried at 60°C and weighed. Roots were collected and scanned as described above. Roots were divided into classes, with nodal roots having a diameter greater than 0.6 mm, thick lateral roots having a diameter greater than 0.3 mm, and thin lateral roots having a diameter less than 0.3 mm.

For the 2019 experiment, four inbred maize genotypes (LH199, PHZ51, DKIB014, and PHG90) were selected based on rooting depth differences in the field in previous experiments. The pots were prepared as follows. Media was placed into the mesocosm to each layer depth, saturated with nutrient solution, and capped with a plastic divider. Dividers were placed at 85, 90, 95, 100, 105, 110, 120, and 130 cm from the top of the column while filling to separate root length by depth. Dividers were made from Plastic Mesh Gutter Guard material (Frost King, Thermwell Products Co., Inc, Mahwah, NJ) so that roots could easily pass through without impedance. The nutrient solution consisted of 2 M ammonium nitrate, 2 M potassium phosphate dibasic, 6 M potassium nitrate, 4 M calcium chloride, 2 M magnesium sulfate, 50 µM potassium chloride, 25 µM boric acid, 2 µM manganese sulfate, 2 µM zinc sulfate, 0.5 µM cupric sulfate, 0.5 µM ammonium molybdate, and 50 µM Fe-DTPA. Nutrient solution pH was maintained at 6.0 with the addition of potassium hydroxide. The nutrient solution applied to the layer from 90-95 cm contained the strontium treatment, in which the calcium chloride in the nutrient solution was replaced with 4 M strontium chloride. When filling the mesocosm, the media of each layer was filled with the same volume and saturated with nutrient solution before placing a divider and filling the next layer. Plants were planted in a staggered manner over 10 days to allow for staggered harvesting, with the first planting occurring on 27 September 2019. Plants received 200 mL of nutrient solution daily for two weeks, and weekly after that through the duration of the experiment. XRF scans were completed on the oldest leaf every three days starting 21 days after germination and continuing until harvest (parameters below). Plants were destructively harvested 28, 38, and 48 days after germination, with three plants of each genotype harvested at each time point. All roots were collected from each media layer, stored in 70% ethanol and later scanned as described previously for length and analysis in WinRHIZO. All roots from the 0-85 cm layer were collected, the number of axial roots per whorl was counted, and one representative root per whorl was scanned. Roots with a diameter less than 0.2 mm were classified as fine roots. After scanning, root samples were oven-dried (60°C) for two days and weighed. Leaves and stem tissue were collected and oven-dried for two days. Leaf discs were collected by taking 1 cm punches from the lamina of the leaf tissue. A homogenous 20 g subsample of each media layer was collected and oven dried for two days prior to elemental analysis.

### 2.4 Field strontium studies

For strontium injection studies, 0.02 moles SrCl_2_ was applied as 10 mL of 2M SrCl_2_ at 30 cm by removing the top 30 cm of soil via a 1.9 cm diameter soil probe. The SrCl_2_ solution was injected using a 30 cm tube with flexible polypropylene tubing onto the soil surface in the hole. The removed soil was then replaced. The injection hole was in-row, between two plants that were later sampled. The site was marked with a flag and plants were tagged for identification. Injection sites were in the genotype DKC49 x 94RIB and adjacent to the plots used in other experiments (2.1 Field LEADER experiments). At six, eight, and ten weeks after germination, plants and soil were sampled. Twelve plants surrounding the Sr injection site were harvested, all leaves and stems collected, air dried, and saved for analysis. Three soil cores around each injection site were collected for soil analysis: one at the point of injection, one in-row, one plant above or below the injection site, and one in a neighboring row between two plants. Four cores were taken for root length analysis, one 5 cm from the base of each plant immediately neighboring the injection site, one 5 cm from the base of the next plant in-row of the injection site, and one 5 cm from the base of the plant in the neighboring row. All cores for root analysis were taken in-row. Cores were processed as described above.

### 2.5 Elemental analysis with XRF

A Bruker Tracer 5i (Bruker Corporation, Billerica, MA, USA) was used for all XRF analysis. For greenhouse experiments on live plants, the oldest green leaf was affixed to the window using micropore tape (3M) placed outside of the view of the analysis window. For all other analysis, dried leaf tissue was placed on the nose of the instrument in a stack of 10, 1 cm discs secured with micropore tape placed outside of the analysis window. Scans were run using different analysis parameters to emphasize different spectral regions as required by the goal of the experiment. Scan parameters for greenhouse experiments and strontium tracer field experiments were 45 kV voltage, 30 µA current, and Cu 38 µm/ Ti 25 µm / Al 300 µm filter for 300 seconds, which emphasized elements in the mid-range of the XRF capabilities, including Sr, Ca, and micronutrients. For other field experiments, we used three additional scans to emphasize different spectral regions. A general scan was first completed at 20 kV, 200 µA, with the Al 38 µm filter. To emphasize light elements, we scanned at 10 kV, 20 µA, and no filter. To obtain counts for heavier elements, we scanned at 30 kV, 30 µA with the Ti 25 µm/Al 300 µm filter. All scans were run for 300 seconds. Counts were extracted in the Artax software (version 8.0.0.443, Bruker) following a Bayesian deconvolution of all elements present in the sample identified manually. Escape and background corrections (9 cycles, 1-40 keV) were applied and data were normalized to the Rh peak. Elemental data was extracted for each element from a single set of scan parameters and a single line from the data set. Details on the extraction of each element from scans can be found in Supplemental Table 2.

For soil analysis, 2g of dried, sieved soil was packed into sample cups (31.5 mm x 34.5 mm Bruker vented cup, part 160.1777). The soil was backed with cotton to ensure close placement on the analysis window. The soil was covered with 4 µm XRF-safe Prolene (Bruker). Soil was analyzed either using the same scan parameters as above for leaf tissue (for Sr analysis only) or using the built-in GeoExploration calibration with a phase-2 oxide calculation.

### 2.6 Elemental analysis with ICP-OES

Dried leaf tissue was weighed, and approximately 0.1 g of tissue was used for ICP-OES (inductively coupled plasma optical emission spectroscopy) analysis. Tissue was digested using a closed-tube digestion system (Wheal et al. 2011). Briefly, tissue was digested overnight in 2 mL of concentrated HNO_3_ and 100 µL of H_2_O_2_ in a closed digestion tube (Starplex Versatube VT502). Prior to digestion 1 mL of HCl was added. Samples were digested for 2 hours at 95°C in a block heater (Environmental Express SC154) and were vented every 30 minutes. Volume was adjusted to 35 mL to a final concentration of 2% HNO_3_ in water and samples were analyzed on a Perkin Elmer Avio 200 ICP-OES equipped with an S10 autosampler.

Optimum wavelengths for each element were determined by comparing with the percent recovery of a known sample (NAPT 2020-206 corn tissue, North American Proficiency Testing, Madison, WI). Wavelengths used were Al 396.154, B 249.677, Ba 233.527, Ca 315.887, Co 228.616, Cr 267,716, Cu 324.752, Fe 238.204, K 766,490, Mg 285.213, Mn 257.610, Na 589.592, N 231.604, P 213.617, Pb 220353, Rb 780.023, S 181.975, Si 251.611, Sr 407.771, Ti 334.940, and Zn 206.200. All peaks were examined manually for background detection and interference using Syngistix for ICP version 3.0 (Perkin Elmer).

For greenhouse and strontium injection field experiments, soil was extracted using an ammonium acetate extraction to extract cations. A 1M ammonium acetate (NH_4_OAc) solution was prepared to a pH of 7.0. Soil (2g) was extracted in a volume of 20 mL by shaking at 150 rpm for 30 minutes in 50 mL conical tubes. After shaking, 10 mL of solution was syringe filtered (0.45 µm polyethersulfone membrane) and 5 mL of 6% HNO_3_ was added prior to ICP analysis. For all other soil analysis, soil was extracted using a Mehlich III protocol prior to ICP analysis (Mehlich 1984).

### 2.7 Data Analysis

All data were analyzed in R version 4.2.2 (R Core Team 2022). Figures were made using ggplot2 3.4.0 (Wickham 2016) with the additional packages of rstatix 0.7.1 (Kassambara 2022), ggh4x 0.2.3 (van den Brand 2022), patchwork 1.1.2 (Pedersen 2022), ggrepel 0.9.2 (Slowikowski 2022), and viridis 0.6.2 (Garnier et al. 2021). Data wrangling was completed using tidyverse 1.3.2 (Wickham et al. 2019). Principal component analyses (PCAs) were completed using centered and scaled data. Clustering was completed using mclust 6.0.0 (Scrucca et al. 2016). Linear discriminant analysis was completed using either a 70-30 training-test set split using the ‘createDataPartition’ function from caret 6.0-93 (Kuhn 2022) or a leave-one-out-cross-validation from the ‘lda’ function from MASS 7.3.58.1 (Venables and Ripley 2002).

## 3. Results

### 3.1 Root depth of field-grown plants correlates with leaf elemental content

We grew four genotypes of maize in the field and measured root depth and collected leaf samples at 6, 8, and 10 weeks after planting. Root depth profiles of the four genotypes show differential growth habits across this time (Figure 1A). The two hybrid lines, DKC25 x 88RIB and DKC49 x 94RIB increased their shallow root development at 8 and 10 weeks of growth, resulting in shallower D95 (the depth at which 95% of the root length has accumulated) values at later time points, which is especially noticeable in DKC35 x 88RIB, the earlier maturing of the two hybrids (88 versus 94 days). In contrast, B73 had an increasing D95 with time, though the values were always less than the other inbred, PHB47, which had its deepest D95 at 8 weeks. This provided us with a range of root depth metrics with which to test our LEADER concept.

**Figure 1.**
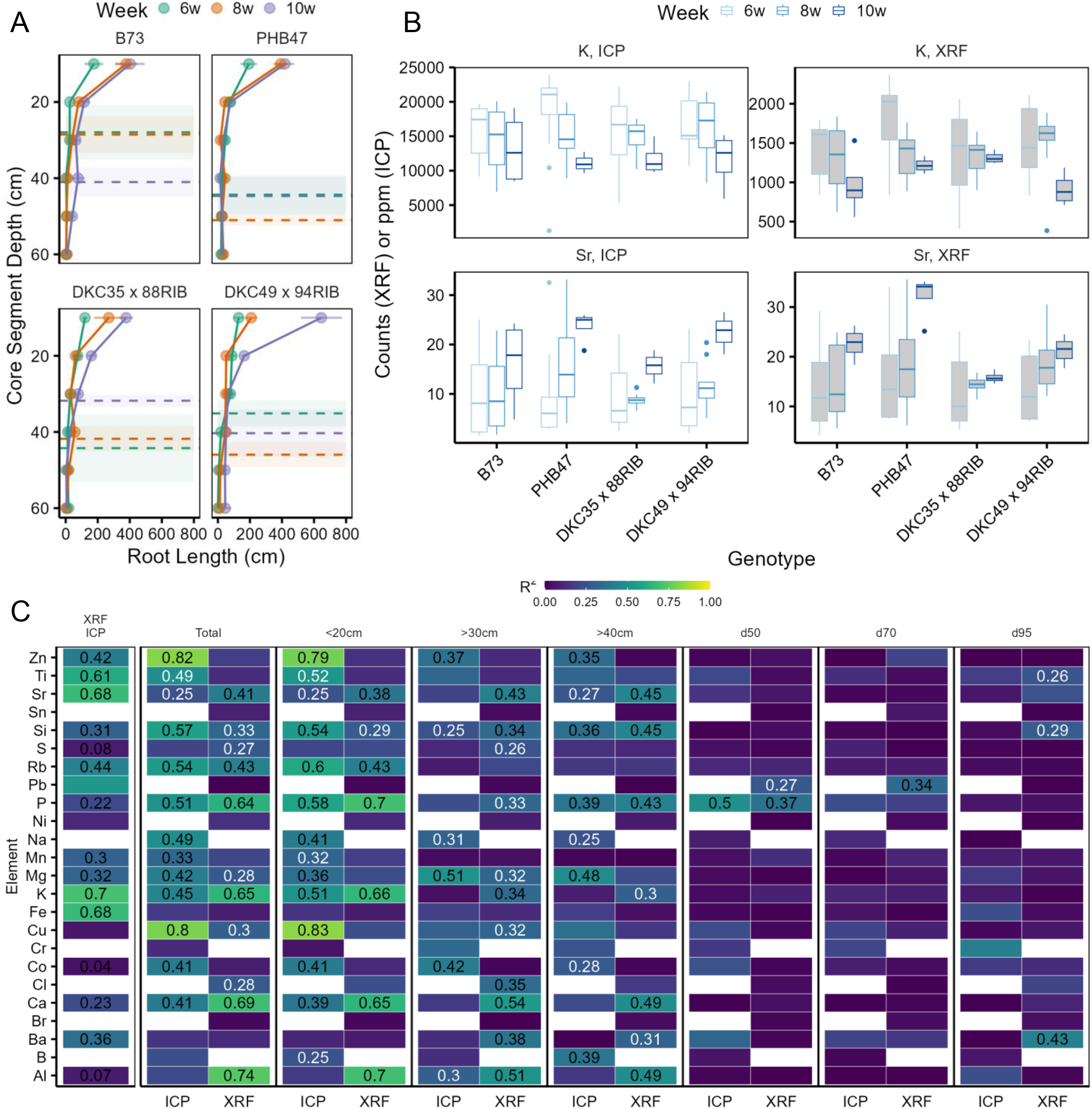
A) Root depth distribution profiles of four maize genotypes. Points represent the root length in each 10 cm layer and error bars represent the standard error of the mean. Dotted lines represent the d95 and shaded regions, the standard error. Color indicates weeks after planting. n = 3 for each datapoint. B) Elemental accumulation of selected elements determined via ICP (unfilled) or XRF (filled) boxplots for young, ear, and old leaves combined. C) Correlations (R^2^) between individual XRF or ICP elements and selected root traits. The first column represents the correlation between XRF and ICP measurements for each element. Subsequent columns represent the relationship between root lengths (total root length, stratified by depth, or D50/D75/D90) and elemental measurements taken as a plot average across all genotypes and time points (n = 12). Missing points are for elements that could not be measured by one of the systems (XRF or ICP). The color of the R^2^ coefficient represents the significance of the relationship, with black numbers indicative of p ≤0.05 and white, p ≤ 0.1.

Leaves are an imperfect subject for quantitative XRF data due to their low density and cellulose matrix. However, we observed similar trends in XRF and ICP data for a number of elements that are measured in both systems (examples shown in Figure 1B). The correlation between XRF counts and ICP ppm values range from R^2^ values of nearly 0 for certain elements that are present in extremely low quantities or are not reliably measurable by one system (e.g. Co, Al, S) to values greater than 0.6 for Ti, Sr, K, and Fe (Figure 1C). Of the 16 elements measured on both systems, 10 correlated with an R^2^ > 0.3 and fifteen were significant at the p ≤ 0.05 level.

Due to the inherent variability in root depth metrics estimated by coring, we took the average values of each plot for both root depth metrics and leaf elemental content metrics. Correlations between single elemental values for ear leaves and roots were not always consistent between ICP and XRF measurements (Figure 1C). ICP data for Zn correlated well with total root length (R^2^ = 0.82, p ≤ 0.05), but XRF data did not show the same relationship (R^2^ = 0.18, ns). However, some elements, including P and K, correlated well with total and shallow root length metrics using both methods of measurement. For deep root metrics, which are more difficult to obtain, XRF data correlated with an R^2^ > 0.3 and p ≤ 0.05 for Sr, Si, P, K, Cu, Cl, Ca, Ba, and Al. ICP data showed a correlation of R^2^ > 0.3 between root length deeper than 30 cm and Zn, Na, Mg, Co, and Al (Figure 1C). Correlations were low for data between individual roots and leaf tissue (Supplemental Figure 1 and 2). We also added soil data into these correlations and found that information about the local soil environment did not improve the correlations (Supplemental Figures 3 and 4). That is, differences in leaf elemental content were not driven by minor differences in elemental content in the soil. A multiple regression between leaf elemental content and the combination of soil elemental content and root depth showed many strong relationships (Supplemental Figures 3 and 4); however, due to the small number of soil samples (n=12), we did not find these models to be convincing.

**Figure 2.**
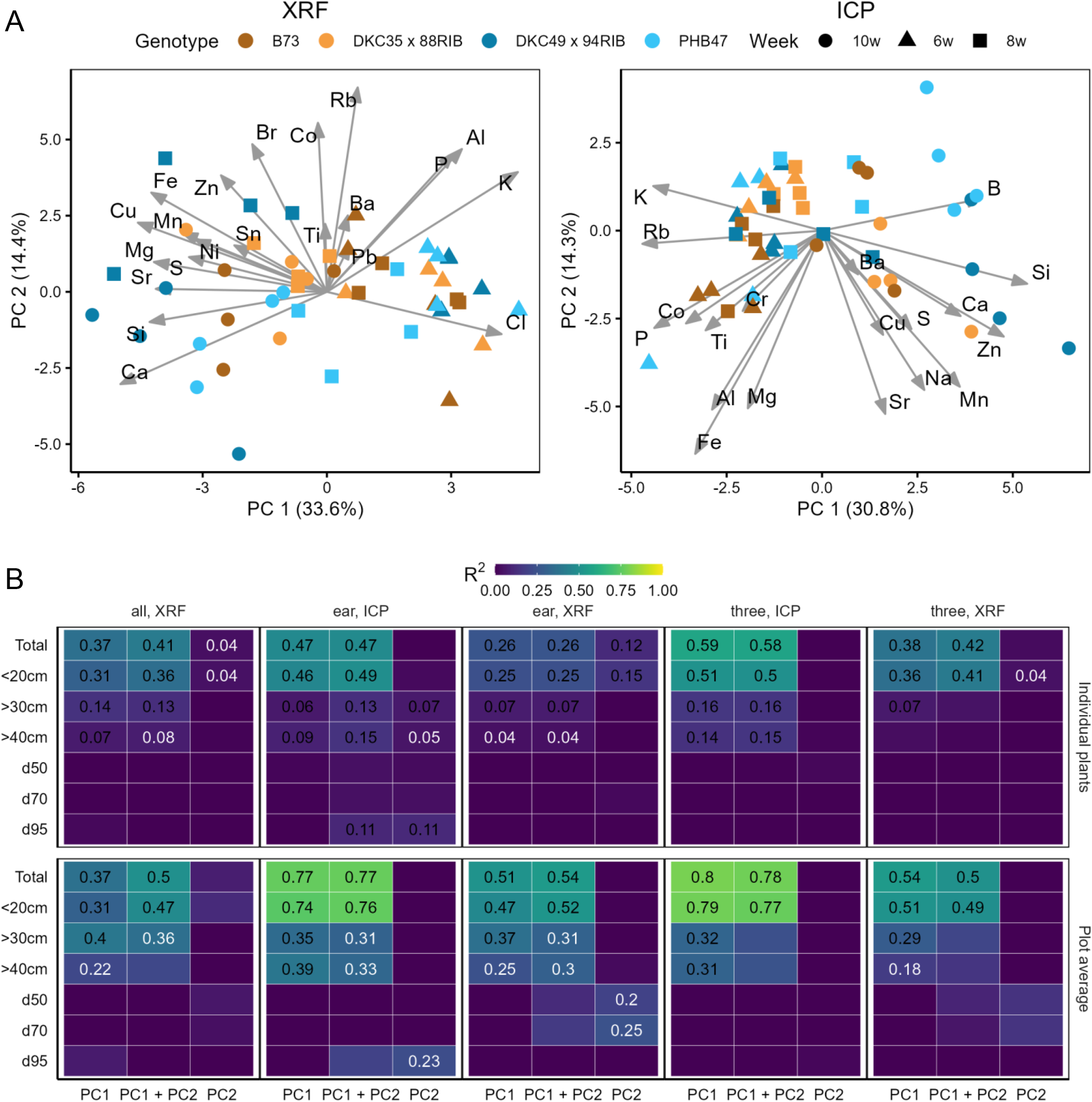
A) PCA biplots of genotypes (color) and timepoints (shape) of leaf elemental content from ear leaves only. Plots represent all individual plants measured via XRF (left) or ICP (right). The first 2 PCs are shown as the X and Y axis with the percent of variation explained by each component included. The loadings of each element are represented with the labeled arrows. B) Adjusted R^2^ values of the relationship between each root trait (Y) with either PC1, PC2, or the combination of PC1 and PC2 measured via XRF or ICP. PCs were constructed using all leaves, only the ear leaf, or a combination of three leaves (oldest, youngest, and ear leaves). Color of the value represents significance at p ≤ 0.05 (black) or p ≤ 0.1 (white).

### 3.2 PCAs correlate with root depth metrics despite noise

The data obtained from either XRF or ICP analysis is highly dimensional, with a single data point having many observations. Since nutrient acquisition and transport within plant tissue results in correlation among individual elements, we employed PCA as a dimension reduction technique to capture information about multiple elements. PCA of XRF and ICP data from individual plant ear leaf tissue (Figure 2A) shows that 1 PC explains 33.6% (XRF) and 30.8% (ICP) of the variation in all elements measured by each system. As expected, elements with similar uptake and/or transport mechanisms, including Ca and Sr, Fe, Cu, and Mn, and K and Rb load similarly on the axes of the PC plots.

We then correlated PCs from each PCA with metrics of root depth (Figure 2B). We used elemental data from individual plants and correlated these with the core taken at the base of an individual plant (top panels) or the plot averages (bottom panels). PCA was performed with either all leaves from each plant (XRF only), the ear leaf only, or a combination of three leaves – the ear, youngest, and oldest leaves from each plant. For ICP measurements, we found strong correlations between PC1 and the combination of PC1 and PC2 for plot average data and ear leaf and combined leaf data, with R^2^ > 0.75 for total and shallow root length metrics. Metrics of deep root length correlated better with plot average data for both XRF and ICP measurements than with data obtained from individual plants, with maximum R^2^ values of 0.4 for root length deeper than 30 cm for plot average data and 0.16 for individual plant data.

### 3.3 Leaf elemental signatures are site dependent

We grew a panel of 30 genotypes at four locations across the US and collected root depth data on a subset of these genotypes (information in Supplemental Table 3). We examined the relationship between elemental content and site by clustering the data (Figure 3A), revealing that clusters are site specific. Of the genotypes sampled, all were found in a minimum of three clusters (Figure 3B). B101, a genotype found in only three clusters, was only grown in Pennsylvania. Genotypes grown at all locations, such as PHZ51 x PHP02, PHG50 x PHG72, PG50, DKPB80 x 3IIH6, DKIB014 x W605D and DKIB014, were found in at least 7 clusters (Figure 3B), indicating that location drove cluster formation more than genotype. Clusters were not specific to treatments, even when treatments were similar (nitrogen and/or water stress) across environments (Figure 3C). Elemental content was largely determined by location (Figure 3D) with clusters being largely specific to an environment with minimal mixtures within a cluster. Due to these differences, individual models to predict or correlate with root depth must be designed for each location independently.

**Figure 3.**
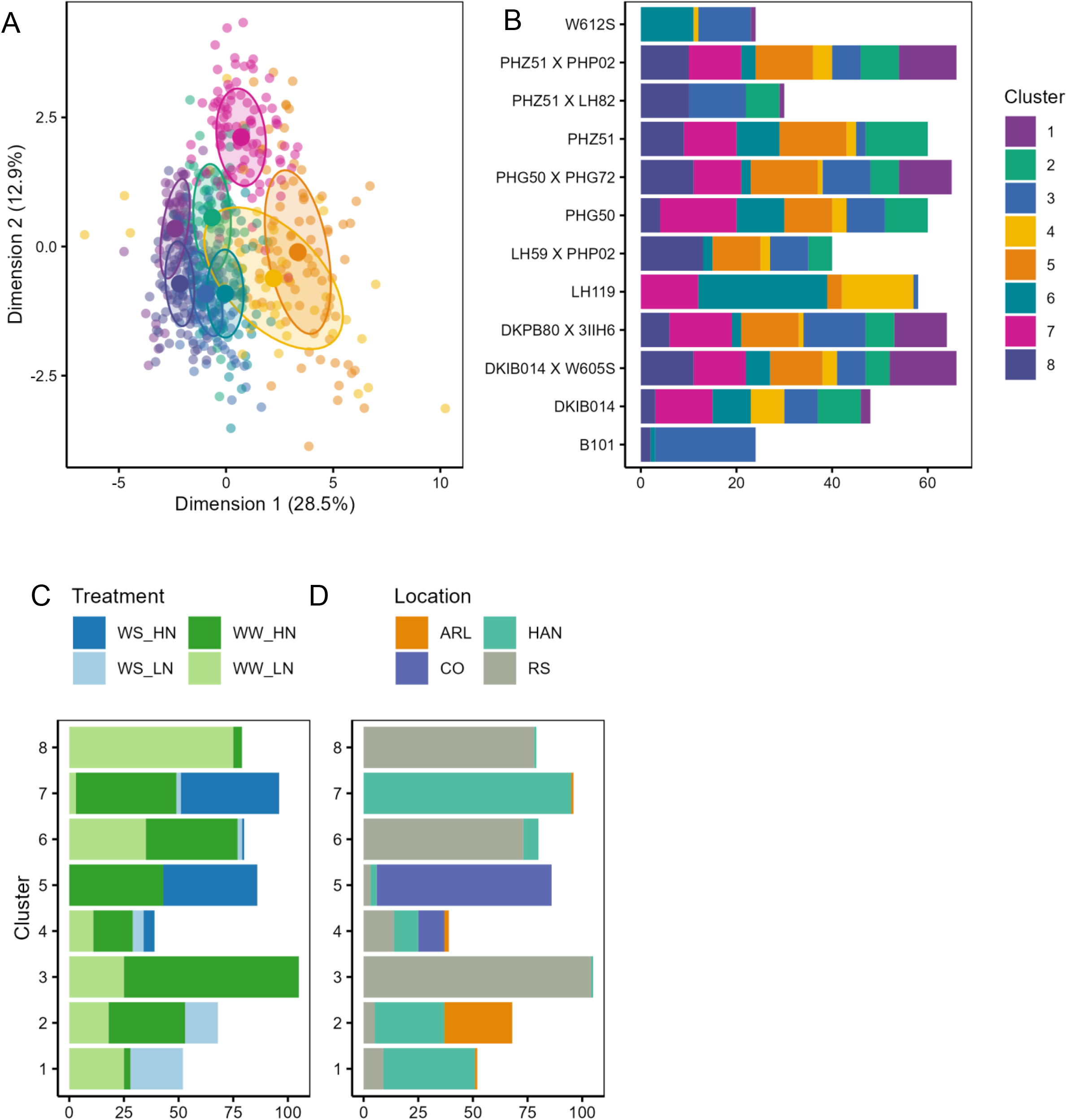
A) Clusters of leaf elemental content determined by the mclust algorithm. Colors represent individual clusters. B) Cluster identity of each genotype. The x-axis represents individuals in each cluster C) Cluster breakdown by treatment. D) Cluster breakdown by experiment location.

### 3.4 Leaf elemental content groups predict root phenotypes in multiple locations

Using the knowledge that elemental content is site-dependent (Figure 3), we examined datasets from four locations (Arlington, WI (ARL), Hancock, WI (HAN), Fort Collins, CO (CO), and Rock Spring, PA (PA)). We used two years of data from PA, with a different set of genotypes grown each year (PA 2019 and PA 2020). Plants grown in PA were subject to a low nitrogen treatment, plants in CO were subject to a water deficit treatment, and plants in Hancock, WI were subject to water deficit, low nitrogen, and the combination of the treatments. We collected XRF data and cores for root length on a subset of genotypes at each location. Information on the genotypes and number of samples taken at each site-year can be found in Supplemental Tables 3 and 4. We applied a similar framework as above, using PCs to predict root traits across these genotypes, but we did not find strong correlations across sites (Supplemental Figures 5-8).

Estimates of root depth from coring are noisy, with variation introduced through small differences in the soil environment, placement of the soil probe, and inherent challenges in the methodology. Rather than relying on specific correlations to explain the relationships between elemental content and root depth, we moved to a classification approach. We used a linear discriminant analysis (LDA) to classify plants at two levels of each of seven root traits: total root length, root length shallower than 20 cm, root length deeper than 30 cm, root length deeper than 40 cm, D50, D80, and D95. We tested these models on a 70/30 training/test split and also employed leave-one-out cross-validation (LOOCV) for additional validation, especially on datasets with a small number of samples.

Elemental content varies with treatment and element, with some elements accumulating more in plants under nitrogen stress, such as Br, some accumulating less in N-stressed plants (Ca), and some showing no difference with treatment (K) (Figure 4A). To account for this, we modeled both a two solution (high and low for each root trait) and a four-solution (high and low for each root trait and each treatment) situation. For each two-solution situation, we classified each plot as either high or low for each root trait by excluding the middle 20% of data centered on the mean value and classifying the top 40% as high and the bottom 40% as low. An example of the trait distribution in a two-solution model for root length deeper than 40 cm the PA 2019 data is in Figure 4B. The data for a four-solution model can be seen in Figure 4C, where the groups are assigned based on the nitrogen treatment and the root length below 40 cm. In this example, there is no significant difference in root length between the nitrogen treatments. This was done for each combination of treatments and root traits.

**Figure 4.**
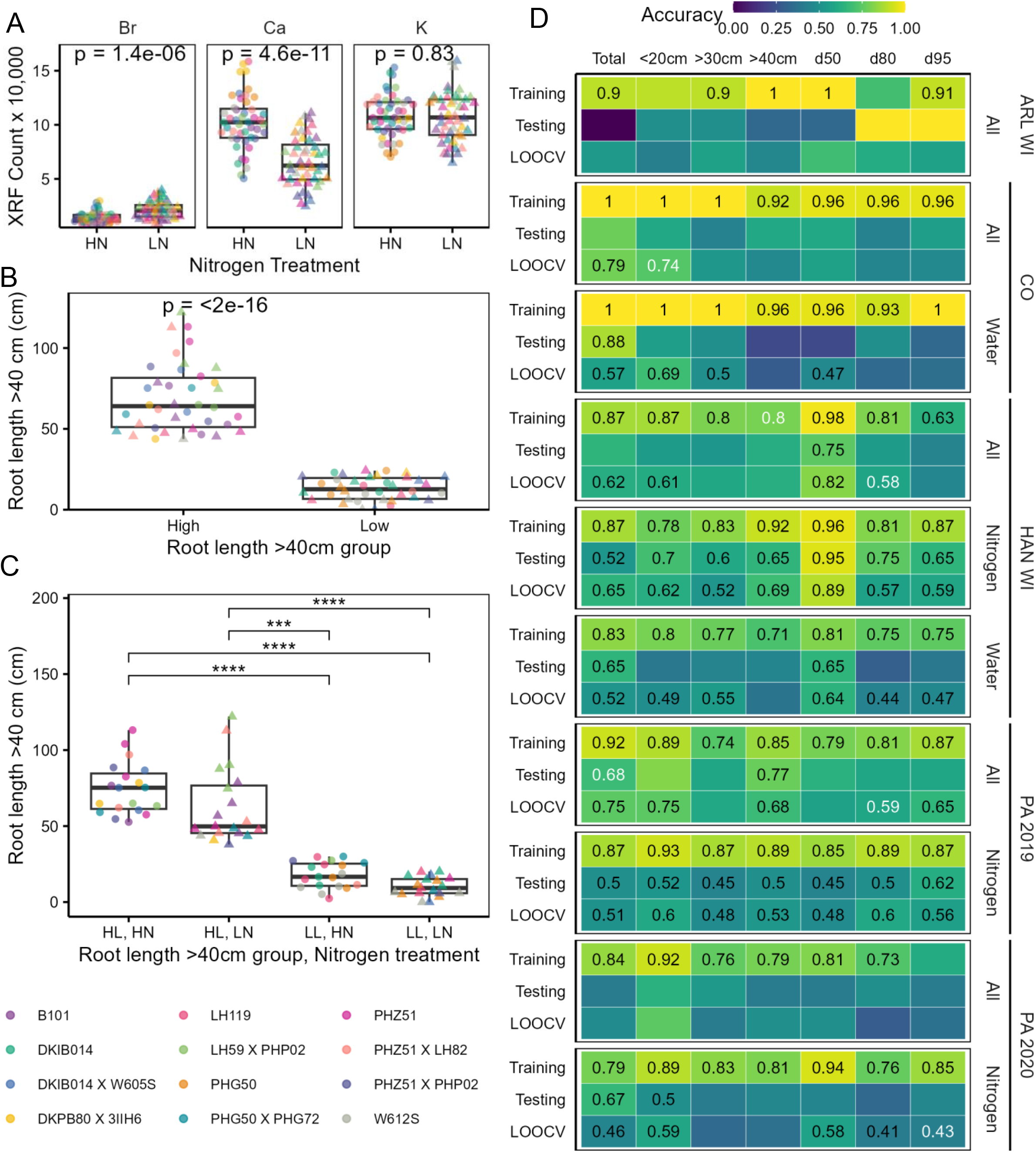
A) Sample elemental content of leaves from plants grown under different nitrogen treatments in PA in 2019 for Br, Ca and K. Displayed p-values are from a t-test. B) Classification of plots into either high or low root length past 40 cm of soil depth. Shapes represent N treatment and colors, genotypes. C) Combinations of N treatment and root length groups from B. Significance from a Dunn test is represented by asterisks, with *** = p ≤ 0.01, and **** = p ≤ 0.001. D) Accuracy values from a LDA of elemental content with root traits across locations. Nitrogen or Water labels indicate a four-solution LDA split by root traits and either N or water treatment. Black values represent significant models at p ≤ 0.05 and white values at p ≤ 0.1. No value indicates no significance of the model.

LDA accuracies for each model (one LD as a predictor) of a set are shown in Figure 4D. Values represent the accuracy of correct class identification and are shown in black for values significant at p ≤ 0.05, white for values significant at p ≤ 0.1, or not displayed for a non-significant (p >0.1) classification. Values are shown for accuracy of assignment for the training data (70%), the testing data (30%), or a LOOCV dataset. For some datasets, sample size was small (ARL WI, n = 18), and performance outside of the training set was poor. The best model performance occurred in our HAN WI and PA 2019 datasets, where we had 2 (HAN WI) or 3 (PA 2019) cores for root depth and leaf samples taken from each plot with 96 plots in each location. This indicates that sample size for training must be large enough to capture variation and allow for robust, significant classification. Mean values were used for each plot to get an accurate representation of plant root traits within a plot. The larger sample size allowed for the development of better models and better classification in both locations. We see improved classification when solving a four-solution model in PA and in HAN WI datasets, with the latter being split based on both nitrogen and water treatment, but not the combination due to the reduction in sample size (Figure 4D).

In the HAN WI dataset, we see robust classification of plots based on the D50 metric across both the two-and-four-solution models. At this site, most root growth occurred above 30 cm (Supplemental Figure 7). As a result, our inability to classify deep root length past 30 or 40 cm in all lines is not surprising. Even with these physical restrictions on root growth, we can classify lines into those with high or low total root length (LOOCV on all lines accuracy of 62%) and shallow root length (LOOCV on all lines accuracy of 61%). In a four-solution model with plots split by nitrogen treatment, our classification accuracy of all root traits is significant, which indicates that we are able to classify nitrogen treatment based upon elemental content as well as the root trait group within that nitrogen class. Separation due to water treatment was not as universally significant, which may partially be due to the water stress treatment in Hancock relying on differential irrigation within natural rainfall. The water-stressed plants received less irrigation, but also received water from rainfall, thus the treatment was not as controlled as the nitrogen stress.

In the PA 2019 dataset, we classified total and deep root length across both the testing and LOOCV sets. In the four-solution model, our classification accuracy is significant for all root traits in both the testing and LOOCV datasets. In the PA 2020 dataset, in which we grew different genotypes in the same location, our classifications were not significant for any testing dataset in the two-solution model. We did have significant classification accuracies in the four-solution model for total and shallow root length in the testing and LOOCV datasets, and the D50, D80, and D95 metrics in the LOOCV dataset. In this dataset, we only had one root core sample and one leaf sample per plot, resulting in increasing noise and reduced accuracy of our input data to build the model.

### 3.5 Strontium uptake correlates with root depth in greenhouse studies

In cases where elemental gradients in field soil may not be sufficient to correlate with root depth, a tracer could be applied to the soil to serve as an indicator of root depth. To test this, we first completed greenhouse studies to identify an appropriate tracer and test the validity of the tracer as a predictor of root depth. In 2018, we tested both Strontium (Sr) and Rubidium (Rb) as tracers. Neither tracer impacted shoot or root growth (Supplemental Figure 9), but since Sr is readily available and more affordable than Rb, we chose Sr as our tracer in a second experiment. Using tall (1.6 m) PVC pots, we applied Sr at a depth of 90 cm and measured Sr accumulation in the oldest leaf of four genotypes over time (Supplemental Figure 10). We began to detect Sr accumulation at 28 days after germination (dag), and harvested plants at 28, 38, and 48 dag.

Strontium was largely concentrated where we applied it (Figure 5A), though there was some downward mobility to deeper layers within the mesocosm. At each sampling time, we collected all roots and weighed those from 0-85 cm and obtained root length of those deeper than 85 cm. Root length shifted downward with time for all genotypes (Figure 5B). At harvest, we collected Sr counts via XRF on all living leaves and correlated Sr levels in either leaf 10 (column 1, Figure 5C) or the total summed counts from leaves 8-12 (column 2, Figure 5C). We found strong correlations at all root depths, with the weakest values related to the shallow root mass above where the Sr was applied (R^2^ = 0.3 and 0.28). When all time points were combined, fine and total root length in the regions where Sr was applied correlated with Sr content (0.48 ≤ R^2^ ≤ 0.58), as did deep root length, with a maximum R^2^ = 0.65 for fine deep root length and Sr content in leaf 10.

**Figure 5.**
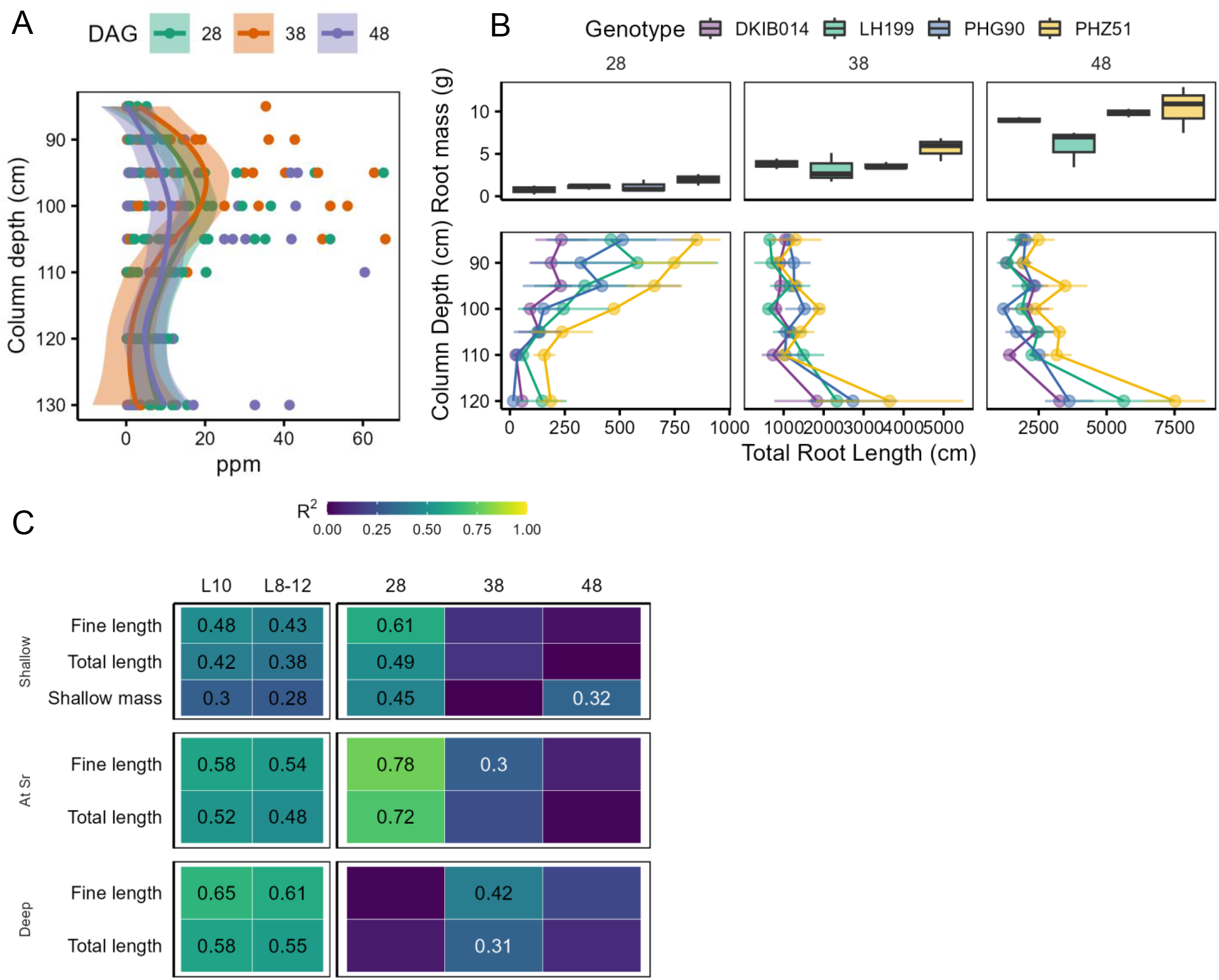
A) Media Sr concentration by depth in greenhouse pots, as determined by ICP. Points represent individual pots colored by harvest date (28, 38, and 48 days after germination). Curves represent the best fit smoothed line. B) Root mass above 85 cm (top panel) and root length distributions below 85 cm (bottom panel) of four genotypes harvested 28, 38, and 48 DAG. C) Correlations between leaf Sr content determined by XRF and root length either above the placement of Sr (95 cm), around the region of Sr (95-105 cm) or below the Sr region. Root length is presented as fine (<0.2 mm) or total root length. Correlations are shown for just leaf 10 (L10) or the summed total of Sr counts in leaves 8-12 in the first two columns. The last three columns represent the summed total from leaves 8-12 at 28, 38, or 48 DAG. Color of the value represents significance at p ≤ 0.05 (black) or p ≤ 0.01 (white).

When leaf Sr from leaves 8-12 was correlated with root depth metrics split by day of harvest, we found significant correlations at the earliest sampling time point at 28 dag between leaf Sr and shallow fine root length (Figure 5C, column 3, R^2^ = 0.61). At 28 dag, there was very little deep root growth, but we did see significant correlations between fine root length (R^2^ = 0.78) and total root length (R^2^ > 0.72, Figure 5B) in the soil layer where Sr was applied. At 38 dag, when deep root growth was present (Figure 5B), we saw significant correlations between leaf Sr and deep fine root length (R^2^ = 0.42, p ≤ 0.05) and deep total root length (R^2^ = 0.31, p ≤ 0.1, Figure 5C, column 4).

### 3.6 Strontium as a tracer in field studies

We applied Sr as a tracer at a depth of 30 cm in field plots of genotype DKC35 x 88RIB. Sr distribution in the soil was measured at 6, 8, and 10 weeks after planting by taking a soil core at the site of application. Sr was largely concentrated in the 20-30 and 30-40 cm layers, with some downward movement in a few samples by 10 weeks (Figure 6A). We sampled plants next to the site of Sr application, within the same row but one plant away from the Sr application, and in the neighboring row. We found plants next to the Sr injection site accumulated significantly more Sr in the older and ear leaves than those in neighboring rows (Figure 6B). Plants within the row of the Sr application accumulated more Sr than those in a neighboring row at 10 weeks in the ear leaf (Figure 6B).

**Figure 6.**
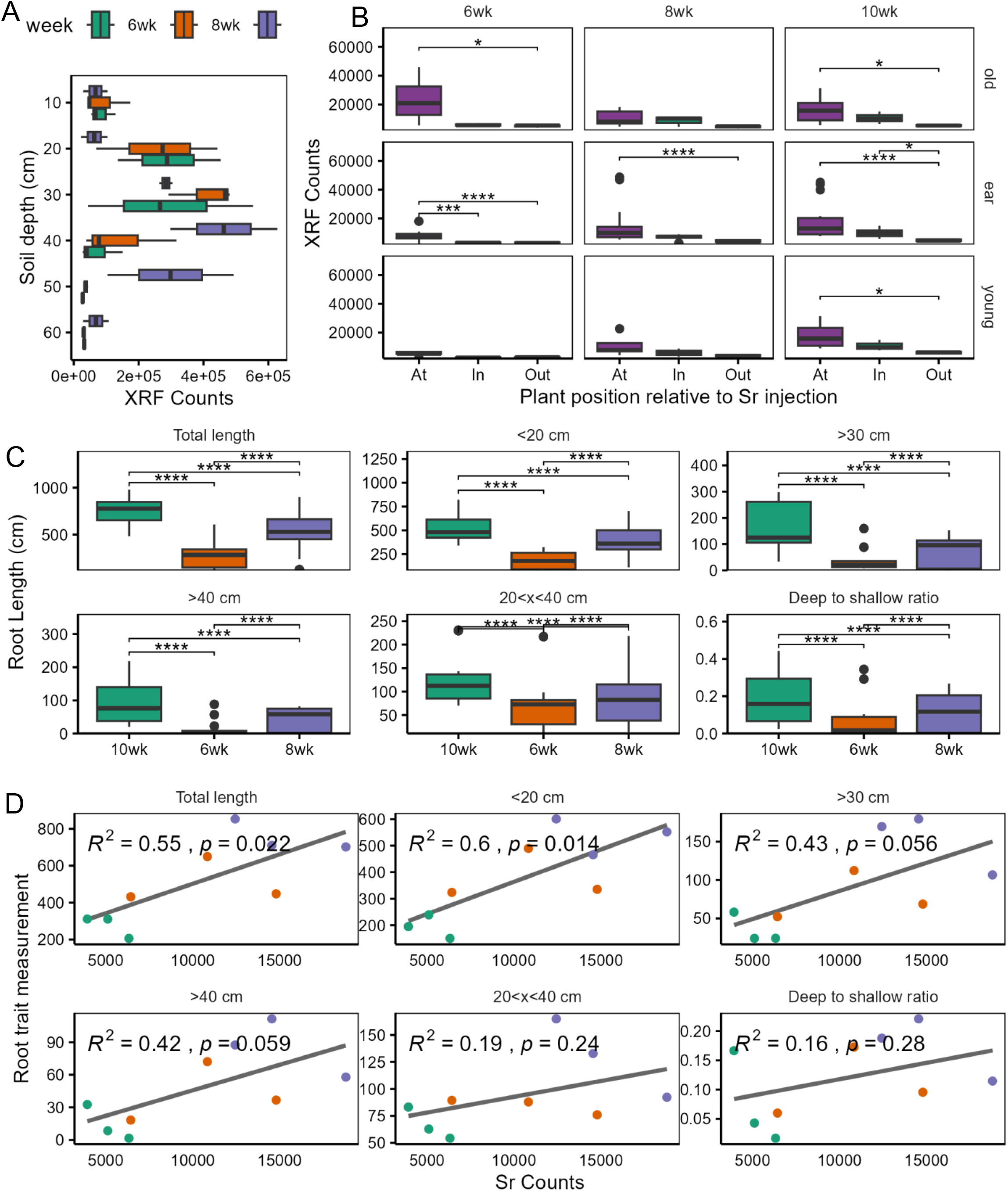
A) Sr content of field soil samples taken where SrCl_2_ was injected into the soil after 6, 8, or 10 weeks determined by XRF. The initial injection was at 30 cm. B) Leaf Sr content determined by XRF of the oldest, ear, and youngest leaves of plants growing at the injection spot, in-row but away from the injection spot (in), or in a neighboring row (out) at 6, 8, and 10 weeks after planting. Significance is from a Dunn test with p ≤ 0.05 (*), p ≤ 0.01 (**), p ≤ 0.001 (***) or p ≤ 0.0001 (****). C) Root length metrics of plants at 6, 8, and 10 weeks after planting. Metrics were taken from 3 cores from each plot with 3 plots sampled at each time point. D) Correlations between root length metrics and leaf Sr content from ear leaves at each time point.

As expected, roots of DKC35 x 88RIB became deeper over time (Figure 6C). We found significant correlations between root length metrics and Sr content in ear leaves for total root length (R^2^= 0.55) and shallow root length (< 20 cm, R^2^ = 0.6). Correlations between deep root length past 30 and 40 cm were significant at the p ≤ 0.1 level (Figure 6D). These results indicate that Sr could be used as a tracer to predict metrics of root depth in soils where natural gradients do not exist or cannot accurately classify or predict root length.

## 4. Discussion

### 4.1 LEADER is a new technology to measure root depth

We present LEADER, a new method to assay root depth of plants growing in the field. Root depth is important for the future of agriculture, with deeper-rooted plants often able to grow better under several edaphic stresses and deeper carbon inputs essential for atmospheric carbon sequestration. An array of root anatomical and architectural phenotypes that increase rooting depth also improve the capture of nitrogen and water (Wasson et al. 2012; Uga et al. 2013; Saengwilai et al. 2014b, a; Chimungu et al. 2014a, 2015; Zhan and Lynch 2015; Zhan et al. 2015; Gao and Lynch 2016; Dathe et al. 2016; Thorup-Kristensen and Kirkegaard 2016; Guo and York 2019; Vanhees et al. 2020, 2022; Wang et al. 2020; Schneider et al. 2022). These phenes are largely related to root number, root angle, root metabolic cost, and the ability of roots to penetrate hard soils, many of which allow the root system to explore soil more efficiently at depth (Lynch et al. 2021; Lynch 2022). The soil environment also shapes root system depth and the ability of roots to grow into deep soil layers (Lynch and Wojciechowski 2015). Low nitrogen and water environments result in deeper rooting (Svoboda and Haberle 2006; Trachsel et al. 2011; Ramamoorthy et al. 2017; Fan et al. 2017). Deep roots are also candidates for increased soil carbon sequestration due to the slower turnover of root-derived carbon at depth (Dietzel et al. 2017), but our ability to leverage deep rooting and our understanding of deep rooting are limited by our ability to measure root depth accurately and easily.

Root depth and distribution change over time and do not increase steadily throughout the season, especially in a crop like maize where new axial root formation starts at or above the soil surface in later developmental stages. Assaying root location is possible through soil coring, minirhizotron imaging, and other technologies introduced previously, but those capture where roots are located at a single moment in time and in space. These tools sample only a small portion of the entire root system, or rhizocanopy (Burridge et al. 2020), which could be an under or overrepresentation of the actual root distribution in soil. Rooting depth has been defined previously using several metrics, including maximum depth and ratio-based calculations, such as D95. Unlike these methods, LEADER relies on the root activity and elemental uptake in its soil environment, which changes with depth in most soils, because of horizonation during pedogenesis, and also because of differential elemental bioavailability with depth associated with gradients in temperature, redox status, pH, water content, and biological activity (Lynch and Wojciechowski, 2015). The accumulation of elements in leaf tissue reflects soil volumes where roots have grown and taken up elements.

Since it is based on basic principles of soil science and plant nutrition, LEADER should work in any plant taxa and in any soil. Here, we showed the utility of LEADER across four distinct regions in the United States with multiple abiotic stress conditions. The soils in these regions were quite different, with only one (Arlington, WI) having a deep Mollisol that is characteristic of the Midwestern corn growing regions. The concepts of LEADER should extend beyond agriculture and would be useful in ecological contexts and in perennial systems. Both herbaceous and woody perennial plants have long-lived roots that can reach soil depths much greater than what is normally seen in agricultural crops (Jackson et al. 1996; Schenk and Jackson 2005; Maeght et al. 2013; Pierret et al. 2016; Thorup-Kristensen et al. 2020; Tumber-Dávila et al. 2022). These plants also grow in non-cultivated soils that should have strong natural elemental gradients. Though tools like soil coring and minirhizotrons have been used in these systems (McCormack et al. 2014; Black et al. 2017; Poeplau et al. 2019; Houde et al. 2020), the depth and extent of roots of perennial plants make coring challenging, and roots of perennial plants could be less active or dead at a time of measurement, making it difficult to determine the active root fraction. Differences in elemental profiles in deep-rooted versus shallow-rooted perennials should be increasingly different from each other with time, which can be explored quickly and repeatedly with LEADER. We also propose that LEADER models should be stable across years in the same fields due to the relationship between soil nutrient availability and soil taxon.

### 4.2 Measuring root depth accurately is challenging

Coring to estimate root depth is an imperfect standard, as are all tools presently in use (Cabal et al. 2021). Various studies have attempted to estimate the number of cores needed to accurately measure root depth in a plot and differentiate genotypes, and these estimates vary from 5 cores per plot (Böhm 2012) to up to 50 cores to reduce relative standard error to 10% (Morandage et al. 2019). Regardless of sampling methods, cores capture all roots within the soil and not only living or active roots. Minirhizotron imaging suffers from the same challenges in addition to problems with presenting an artifactual growth surface that contrasts with the soil. New imaging techniques exploiting multispectral imaging aims to identify and classify roots as living or dead (Svane et al. 2019a), though inherent challenges with relating minirhizotron imaging to true root mass remain (Samson and Sinclair 1994).

The use of tracers allows us to assay where roots are actively capturing resources in soil. Careful tracer application in the semifield RadiMax facility, coupled with minirhizotron imaging, shows that there is potential for careful use of ^15^N as a tracer for measuring root depth, but whether, and how, this scales to field phenotyping still remains to be discovered (Svane et al. 2019b; Chen et al. 2019; Wacker et al. 2022). The use of tracers in rhizobox studies shows that deep root length and tracer uptake are related (Rasmussen et al. 2020), though the utility of this in the field has not been scaled beyond smaller proof-of-concept style work (Hoekstra et al. 2014). Many field studies have used ^15^N to determine root N acquisition, but few have compared ^15^N accumulation to detailed measurements of root depth (Saengwilai et al. 2014b). Here, we used Sr as a tracer in both greenhouse and field studies and found strong correlations with root growth, especially in greenhouse studies. Though the use of Sr as a tracer is not new (Hoekstra et al. 2014; Rasmussen et al. 2020; Han et al. 2020) our ability to measure Sr content over time with XRF allows for the tracing of leaf accumulation non-destructively and quickly, with results available in a matter of minutes. Compared to analyzing tissue using ICP-OES or ICP-MS or measurement of isotopes, XRF is a much faster method that will allow researchers to better understand root placement and activity quickly.

Models for validating new methods for measuring root depth are built assuming that coring or minirhizotron imaging are the ground truth or gold standard for root measurement. We chose to use coring as it was the best and most accessible method, even given its limitations. LEADER, as do all studies using tracers, relies on root activity, uptake, and transport of elements from soil into leaf tissue. As LEADER relies largely on natural elemental gradients in soil, we do not capture a short moment in time as do some studies with applied tracers. Rather, LEADER reflects the residence and activity time of each root in soil layers that contain the elements of interest. In the context of breeding and selecting deep-rooted lines, LEADER presents the possibility of quickly and easily categorizing field-grown plants as deep or shallow-rooted lines. In the context of understanding plant-soil relationships and root growth, LEADER will allow us to non-destructively estimate root depth over time and further disentangle the relationship between a root and its natural soil environment. Understanding the spatiotemporal development of roots in soil is hindered by the soil itself, but information gleaned from LEADER studies, combined with some excavation (coring) and *in silico* studies (Burridge et al. 2020) will allow us to develop a more complete picture of the functioning of an entire root system.

Scaling any technology to measure roots at depth is challenging. LEADER, once validated through coring or another method, should scale quickly and cost-effectively to approximate root depth. Because soil elemental content differs by region, characterization of root depth characteristics is required for each site. Here, we show that LEADER successfully classified root systems by total, deep, and shallow root length at four different sites across the United States. However, our ability to correctly classify root depth was constrained by our sampling size, with the accuracy of models categorizing root depth decreased with small sample size. Recommendations on how many validation cores are needed will vary with each site and the heterogeneity of the soil at that site. Our predictive ability was best when we had a large number of individual cores (n = 192 and n = 288) taken from a large number of individual plots (n = 96). Our ability to categorize root systems into different depth classes decreased with smaller sample sizes (n = 18) or when we sampled with less intensity in a single plot (PA20). We recommend a minimum of 3 cores per plot to estimate root growth metrics, though more cores would result in better models.

### 4.3 Elemental analysis of leaf tissue can be accomplished with XRF

LEADER relies on measuring the elemental content of leaf tissue. We used hand-held XRF spectroscopy as a tool for analysis and found it comparable to the standard ICP-OES system. ICP-OES is widely available, affordable, and accessible to many research groups. ICP-MS (inductively coupled plasma mass spectroscopy) is more precise and can measure more elements at lower concentrations but is more costly and may be less accessible than ICP-OES. Though the standard, ICP-OES suffers from challenges with measuring certain elements that are present in plant tissue, including Cl, Br, and S. The cost of consumables, including reagents for digestion, standards, and gas for running the instrument are not trivial. Though samples can be prepared in batches and run on the instrument using an autosampler, sample processing and analysis takes up to 24 hours.

XRF is not a new technology, with the first handheld, portable systems introduced in 1982, but its use in agricultural sciences is relatively new (Sapkota et al. 2019; Montanha et al. 2020; Khan et al. 2022). The cellulosic matrix, high water content, and thinness of leaf tissue make it an imperfect system for XRF analysis, as not all X-Rays put into the system will be accurately reflected to the detector. XRF with plant tissue, therefore, is difficult to calibrate for a fully quantitative analysis. Here, we use normalized counts to compare plant tissue samples rather than converting these counts to elemental content (ppm) in order to avoid challenges with imperfect calibration. The XRF instrument that we used, Bruker Tracer 5i, can theoretically measure elements from Na to U on the periodic table, but the ability to accurately measure elements lighter than Ar is confounded by air between the detector and the sample, which can be improved with a helium purge, though that decreases throughput and portability. With XRF, there are no consumables, just a thin plastic (Ultralene) window that requires occasional replacement. Furthermore, the time to prepare a sample is minimal and the time to analyze a sample is measured in minutes, not hours. We found correlations between XRF counts and ICP-OES analysis for a number of elements, and these correlations were strong for elements that both platforms are good at measuring. XRF gave us a broader range of elements, including Sn, Pb, Ni, Cl, and Br, that we did not get with our ICP-OES setup. XRF was missing Na, Cr, and B, though Na and Cr were detectable only at low levels using ICP-OES. We completed our analyses using the instrument in the lab on collected leaf discs due to university regulations on X-ray producing devices and their users. However, using handheld XRF in the field, which is already done in soil science and geoscience, would make elemental analysis of plant tissue much faster. The device could be set up at or near the field site, and samples could be analyzed in minutes after collection. This could allow for estimates of root depth using the concepts of LEADER to be calculated quickly and at scale.

### 4.4 Conclusions

We have developed LEADER as a method to estimate root depth of maize in field conditions. We show that leaf elemental profiles differ with rooting depth at multiple field locations across the US, and that these elemental profiles can be used to classify root systems based on their total length or length by depth. We show that Sr can be used as a tracer in both greenhouse and field studies, and that Sr can be detected and measured using XRF. As a tool, LEADER has the potential to increase our understanding of rooting depth, a notoriously difficult aspect of plant and root biology to measure. The concepts of LEADER should be adaptable to other agricultural and natural systems, and we encourage its use for estimating rooting depth.

## Core ideas (3-5 impact statements, 115 char max for each)

1. Root depth of field-grown plants correlates with leaf elemental content
2. Leaf elemental signatures are site dependent
3. Leaf elemental content groups predict root phenotypes in multiple locations
4. Strontium is suitable as an applied elemental tracer of root depth
5. XRF is suitable for in field measurements of leaf elemental content

## Abbreviations

LEADER: Leaf Element Accumulation from Deep Roots
XRF: X-Ray Fluorescence
D95: the depth at which 95% of the root length has accumulated
ICP-AES: Inductively Coupled Plasma Optical Emission spectroscopy

## Supplemental Material

Supplemental Figure 1 Correlations between individual elements obtained from XRF (left column) or ICP (right column) and root traits. Values were obtained from leaves on a single plant, and root trait values were calculated based on root metrics from a single core taken at the base of the same plant. Color of the value represents significance at p ≤ 0.05 (black) or p ≤ 0.01 (white).

Supplemental Figure 2. Correlations between individual elements obtained from XRF (left column) or ICP (right column) and root traits. Values are the average values for a single plot with four plants for leaf elemental values and four cores taken to estimate root metrics. Color of the value represents significance at p ≤ 0.05 (black) or p ≤ 0.01 (white).

Supplemental Figure 3. A) Correlations between total leaf elemental content and soil elemental content as determined by XRF. Soil data were divided to represent shallow (0-20 cm), mid-depth (20-40 cm), deep (40-60 cm) and total (0-60 cm). Elemental data was averaged across an entire plot. B) Adjusted R^2^ values for the linear model for leaf elemental content as predicted by the combination of root length in each depth zone and the respective soil elemental content in the corresponding depth zone determined by XRF. Color of the value represents significance at p ≤ 0.05 (black) or p ≤ 0.01 (white).

Supplemental Figure 4. A) Correlations between total leaf elemental content and soil elemental content as determined by ICP. Soil data were divided to represent shallow (0-20 cm), mid-depth (20-40 cm), deep (40-60 cm) and total (0-60 cm). Elemental data was averaged across an entire plot. B) Adjusted R^2^ values for the linear model for leaf elemental content as predicted by the combination of root length in each depth zone and the respective soil elemental content in the corresponding depth zone determined by ICP. Color of the value represents significance at p ≤ 0.05 (black) or p ≤ 0.01 (white).

Supplemental Figure 5: PA 2019 A) Root depth distributions of 12 genotypes grown in the field under high (HN) or low (LN) nitrogen. The dotted lines and triangles represent the LN treatment. Points represent the root length in each 10 cm layer and error bars represent the standard error of the mean. Dotted lines represent the D95 and shaded regions, the standard error. Color indicates weeks after planting. n = 12 for each datapoint. B) PCA biplots of all plants with triangles representing the LN treatment. Loadings on the PCs are indicated with labeled arrows. C) PCA biplot of mean elemental values of the treatments and genotypes for each treatment group. D) Correlations (R^2^) between root traits (Y) and PC1 of either all individual plots or the mean of elemental values and root values. Correlation values are shown when they are significant at the p ≤ 0.05 level.

Supplemental Figure 6: CO 2019 A) Root depth distributions of 8 genotypes grown in the field under well-watered (WW) or water-stressed (WS) treatments. The dotted lines and triangles represent the WW treatment. Points represent the root length in each 10 cm layer and error bars represent the standard error of the mean. Dotted lines represent the D95 and shaded regions, the standard error. Color indicates weeks after planting. B) PCA biplots of all plants with triangles representing the WW treatment. Loadings on the PCs are indicated with labeled arrows. C) PCA biplot of mean elemental values of the treatments and genotypes for each treatment group. D) Correlations (R^2^) between root traits (Y) and PC1 of either all individual plots or the mean of elemental values and root values. Correlation values are shown when they are significant at the p ≤ 0.05 level.

Supplemental Figure 7: HAN 2019 A) Root depth distributions of 8 genotypes grown in the field under well-watered (WW), water-stressed (WS), high-nitrogen (HN), and low-nitrogen (LN) treatment combinations. Circles represent WS_HN; triangles, WS_LN; squares, WW_HN; and crosses, WW_LN. Points represent the root length in each 10 cm layer and error bars represent the standard error of the mean. Dotted lines represent the D95 and shaded regions, the standard error. Color indicates weeks after planting. B) PCA biplots of all plants with triangles representing the WW treatment. Loadings on the PCs are indicated with labeled arrows. C) PCA biplot of mean elemental values of the treatments and genotypes for each treatment group. D) Correlations (R^2^) between root traits (Y) and PC1 of either all individual plots or the mean of elemental values and root values. Correlation values are shown when they are significant at the p ≤ 0.05 level in black and in white when p ≤ 0.1.

Supplemental Figure 8: ARL 2019 A) Root depth distributions of 6 genotypes grown in the field in Arlington, WI. Points represent the root length in each 10 cm layer and error bars represent the standard error of the mean. Dotted lines represent the D95 and shaded regions, the standard error. Color indicates weeks after planting. B) PCA biplots of all plants with triangles representing the WW treatment. Loadings on the PCs are indicated with labeled arrows. C) PCA biplot of mean elemental values of the treatments and genotypes for each treatment group. D) Correlations (R^2^) between root traits (Y) and PC1 of either all individual plots or the mean of elemental values and root values. Correlation values are shown when they are significant at the p≤ 0.05 level.

Supplemental Figure 9. A) Shoot dry biomass of greenhouse-grown maize plants after 48 days. Tracers for root depth (0.06M Sr – 0.17M Rb) did not impact biomass (ns, Dunn Test). B) Root length metrics of 48-day old greenhouse grown maize plants grown with tracers. There were no significant effects of tracers. C) XRF counts of the oldest, green leaf of plants with tracers applied. Only after 28 days were plants provided 0.34 M Sr showing significant accumulation, though we saw dose-dependent increasing trends (p ≤ 0.05, t-test compared to control conditions).

Supplemental Figure 10. Accumulation of Sr measured by XRF in the oldest green leaf over time in the greenhouse (2019 experiment). Four genotypes were measured (n = 9).

Supplemental Table 1. Agronomic and field information for field sites used in this study

Supplemental Table 2. Element extraction parameters from XRF scans.

Supplemental Table 3. Germplasm used in field studies. X indicates the plants were grown and sampled for leaf elemental content. Xc indicates cores were taken for root depth.

Supplemental Table 4. Sampling information for field studies

## Data Availability

Data and code are available at https://doi.org/10.5281/zenodo.7877682

## Author Contributions

MTH participated in data curation, formal analysis, investigation, methodology, project administration, supervision, visualization, writing (original draft and review and editing). KBE participated in supervision and writing (review and editing). JPL was responsible for conceptualization, funding acquisition, project administration, supervision, methodology, and writing (original draft, review and editing).

## Acknowledgements

We thank Michael Williams and Peter Ilhardt for technical assistance.

