## Supplemental Figures for "LEADER (Leaf Element Accumulation from Deep Roots): a nondestructive phenotyping platform to estimate rooting depth in the field"

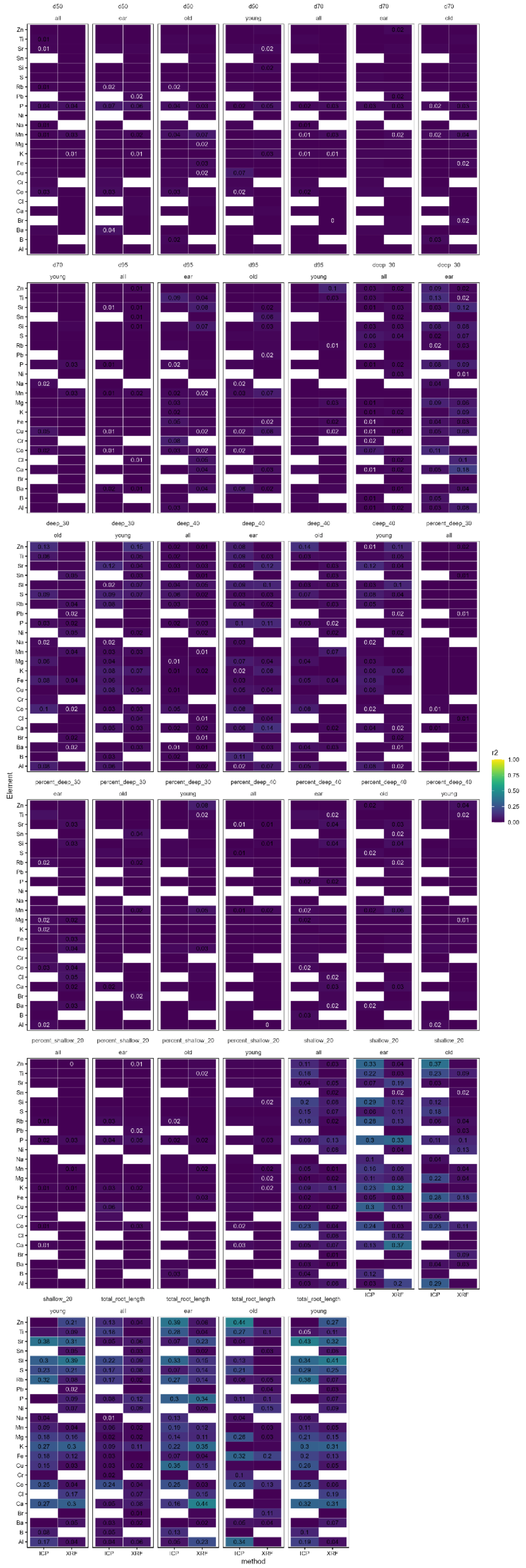

Supplemental Figure 1. Correlations between individual elements obtained from XRF (left column) or ICP (right column) and root traits. Values were obtained from leaves on a single plant, and root trait values were calculated based on root metrics from a single core taken at the base of the same plant. Color of the value represents significance at p ≤ 0.05 (black) or p ≤ 0.01 (white).

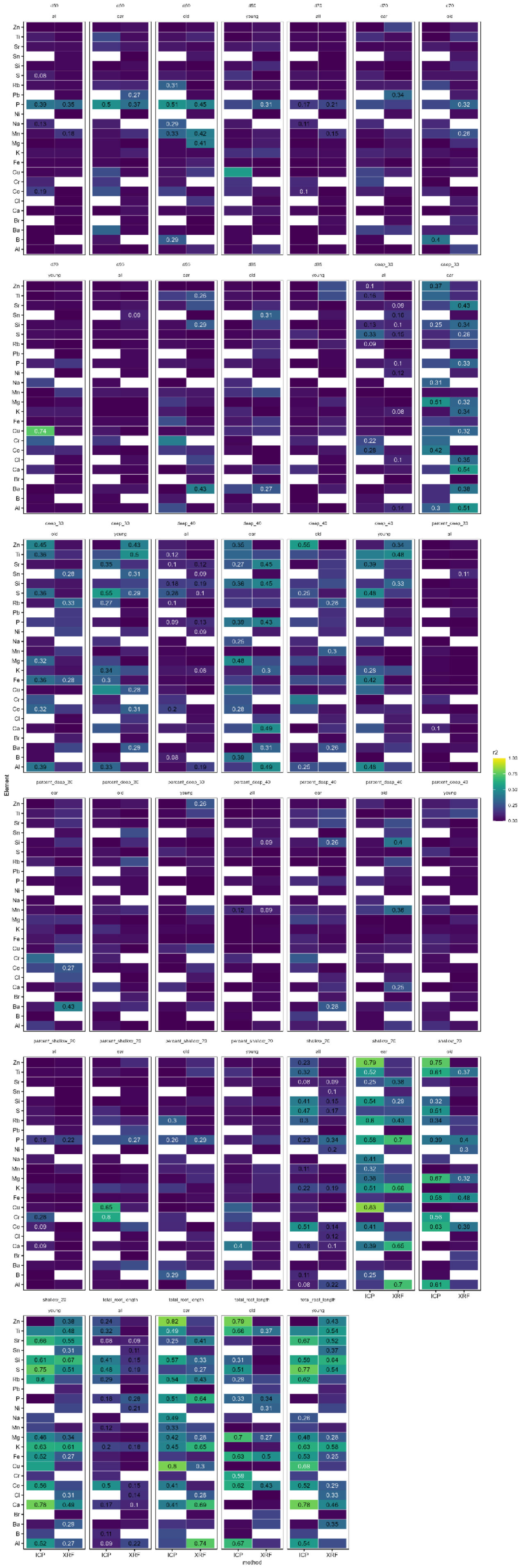

Supplemental Figure 2. Correlations between individual elements obtained from XRF (left column) or ICP (right column) and root traits. Values are the average values for a single plot with four plants for leaf elemental values and four cores taken to estimate root metrics. Color of the value represents significance at p ≤ 0.05 (black) or p ≤ 0.01 (white).

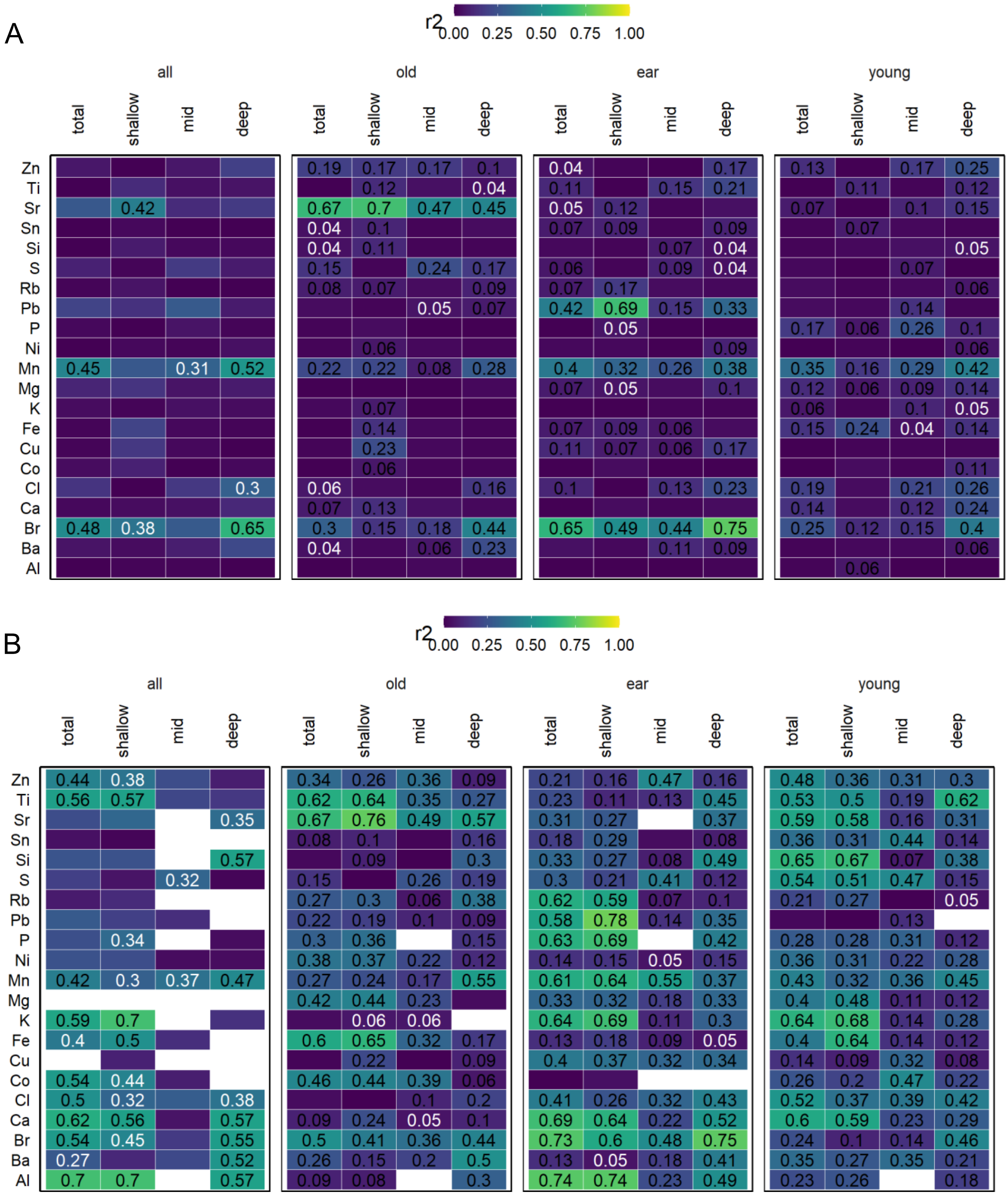


Supplemental Figure 3. A) Correlations between total leaf elemental content and soil elemental content as determined by XRF. Soil data were divided to represent shallow (0-20 cm), mid-depth (20-40 cm), deep (40-60 cm) and total (0-60 cm). Elemental data was averaged across an entire plot. B) Adjusted $R^2$ values for the linear model for leaf elemental content as predicted by the combination of root length in each depth zone and the respective soil elemental content in the corresponding depth zone determined by XRF. Color of the value represents significance at $p \leq 0.05$ (black) or $p \leq 0.01$ (white).

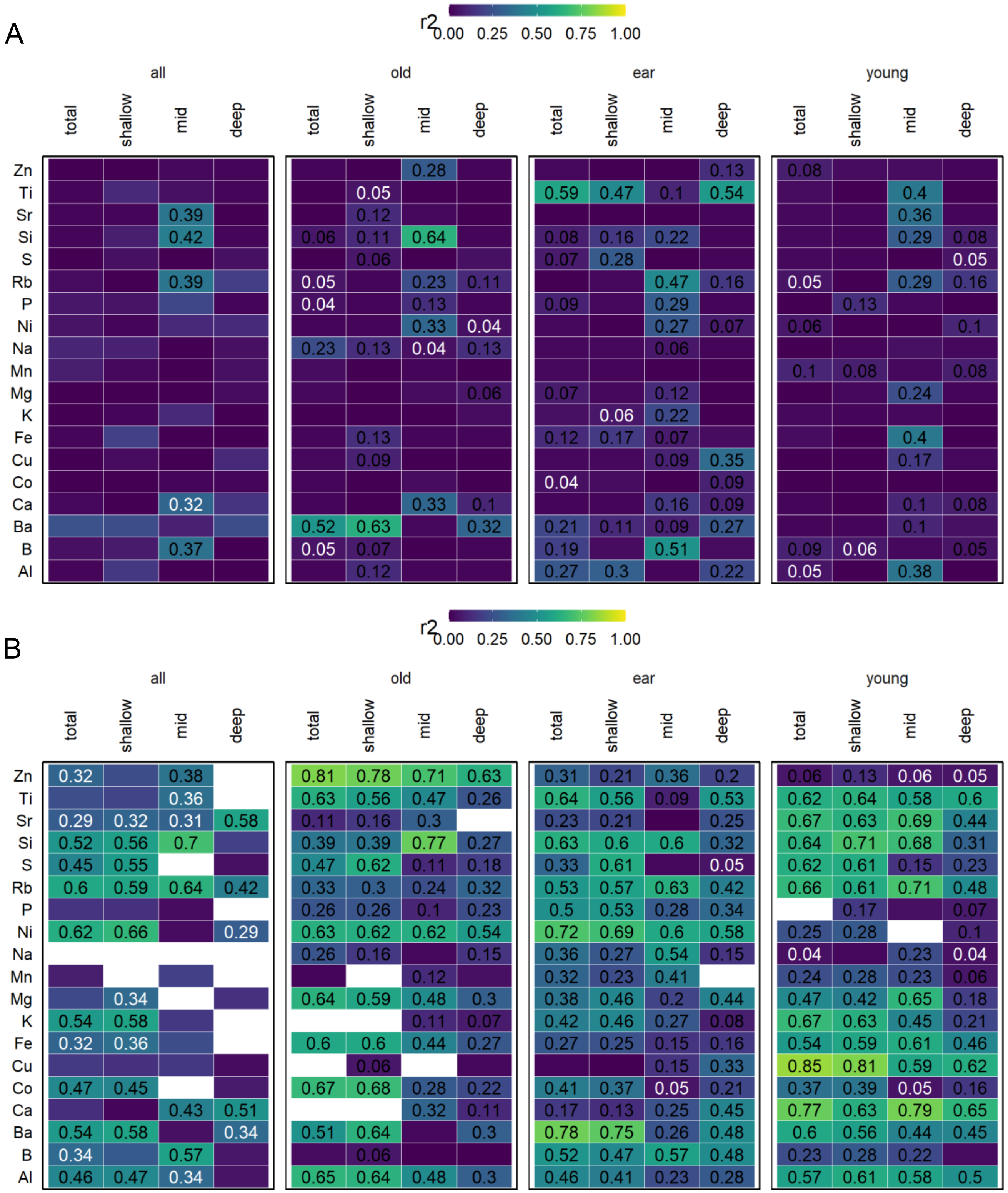


Supplemental Figure 4. A) Correlations between total leaf elemental content and soil elemental content as determined by ICP. Soil data were divided to represent shallow (0-20 cm), mid-depth (20-40 cm), deep (40-60 cm) and total (0-60 cm). Elemental data was averaged across an entire plot. B) Adjusted $R^2$ values for the linear model for leaf elemental content as predicted by the combination of root length in each depth zone and the respective soil elemental content in the corresponding depth zone determined by ICP. Color of the value represents significance at $p \leq 0.05$ (black) or $p \leq 0.01$ (white).

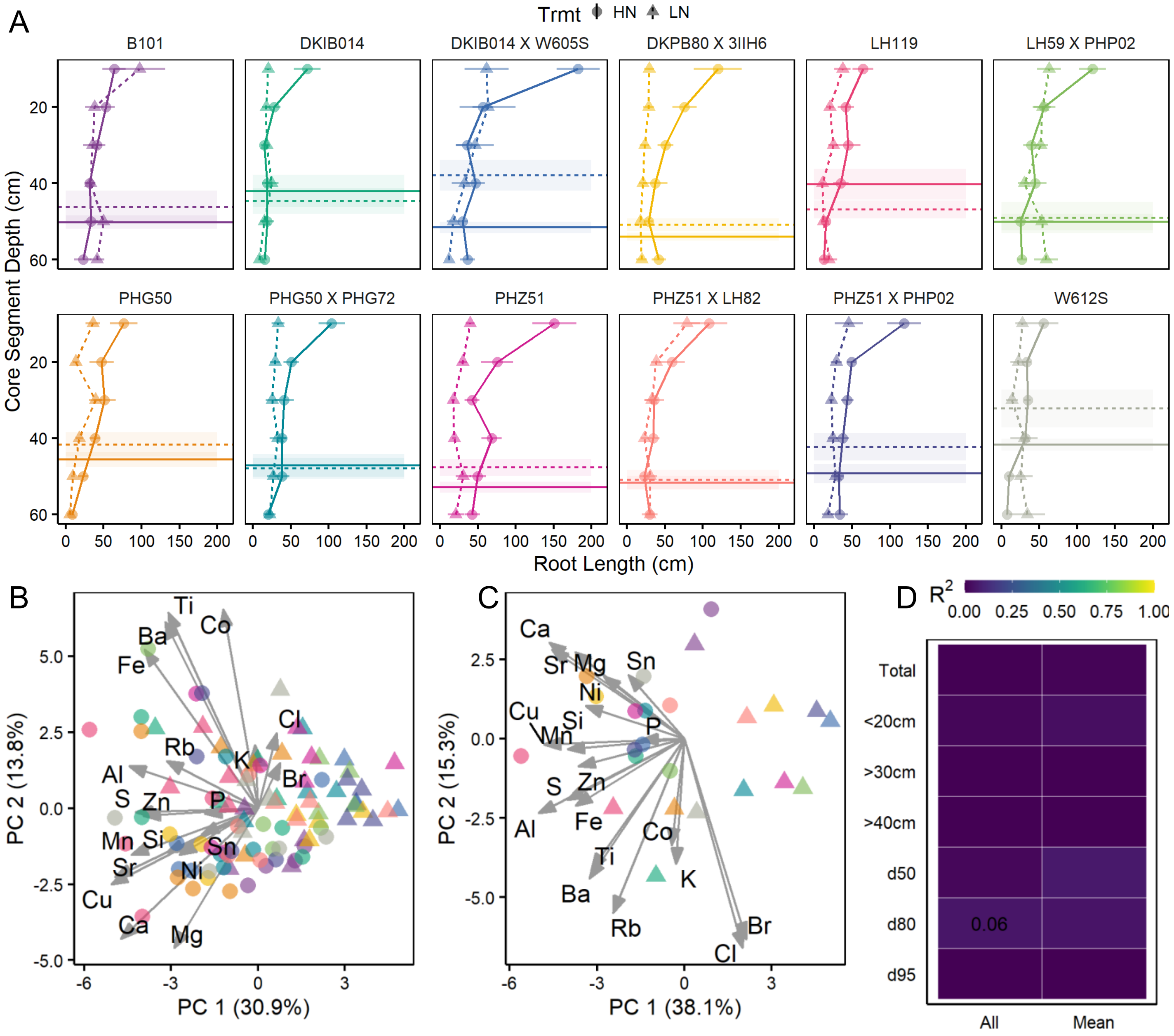


Supplemental Figure 5: PA 2019 A) Root depth distributions of 12 genotypes grown in the field under high (HN) or low (LN) nitrogen. The dotted lines and triangles represent the LN treatment. Points represent the root length in each 10 cm layer and error bars represent the standard error of the mean. Dotted lines represent the D95 and shaded regions, the standard error. Color indicates weeks after planting. n = 12 for each datapoint. B) PCA biplots of all plants with triangles representing the LN treatment. Loadings on the PCs are indicated with labeled arrows. C) PCA biplot of mean elemental values of the treatments and genotypes for each treatment group. D) Correlations ($R^2$) between root traits (Y) and PC1 of either all individual plots or the mean of elemental values and root values. Correlation values are shown when they are significant at the $p \leq 0.05$ level.

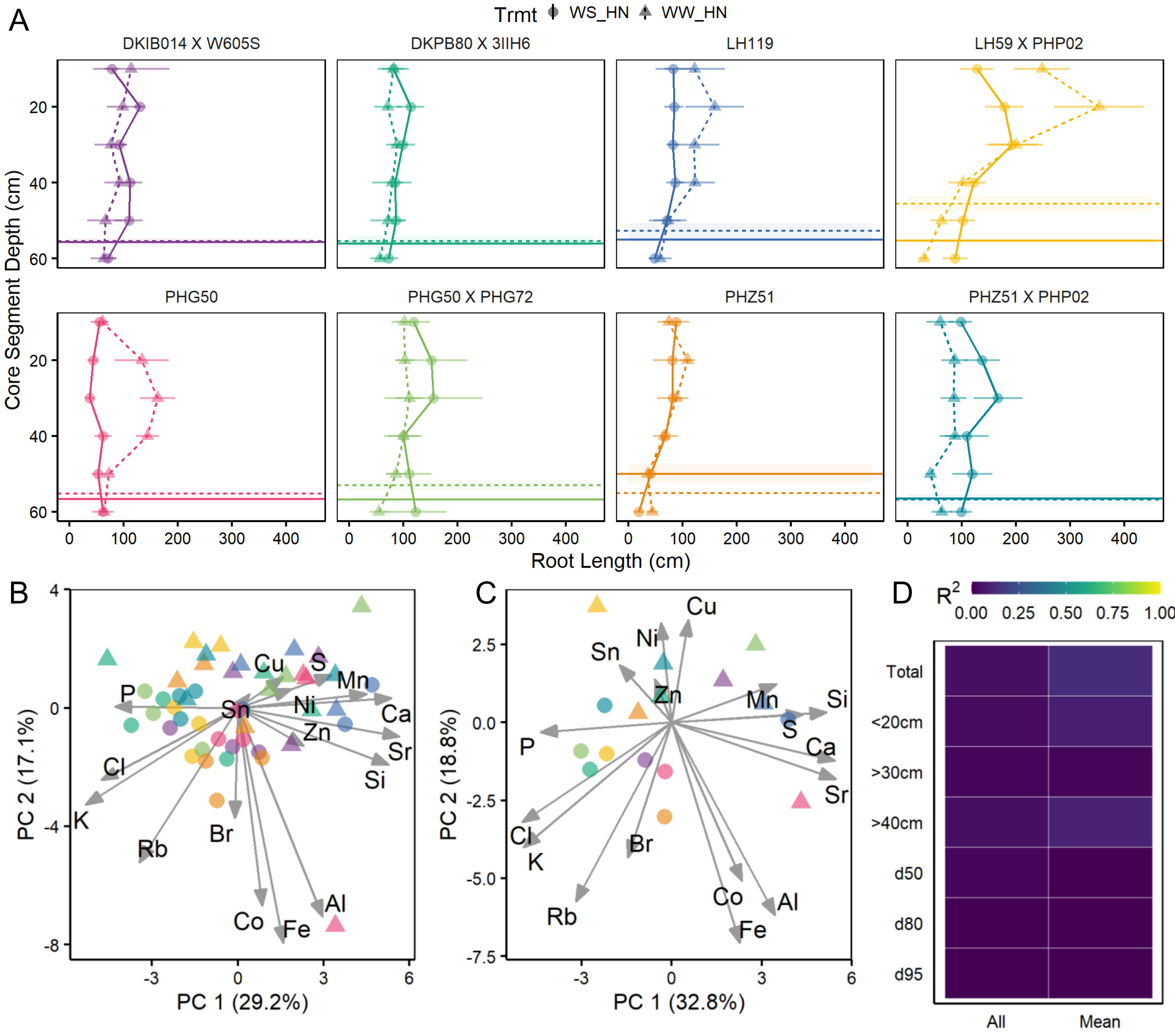


Supplemental Figure 6: CO 2019 A) Root depth distributions of 8 genotypes grown in the field under well-watered (WW) or water-stressed (WS) treatments. The dotted lines and triangles represent the WW treatment. Points represent the root length in each 10 cm layer and error bars represent the standard error of the mean. Dotted lines represent the D95 and shaded regions, the standard error. Color indicates weeks after planting. B) PCA biplots of all plants with triangles representing the WW treatment. Loadings on the PCs are indicated with labeled arrows. C) PCA biplot of mean elemental values of the treatments and genotypes for each treatment group. D) Correlations ($R^2$) between root traits (Y) and PC1 of either all individual plots or the mean of elemental values and root values. Correlation values are shown when they are significant at the $p \leq 0.05$ level.

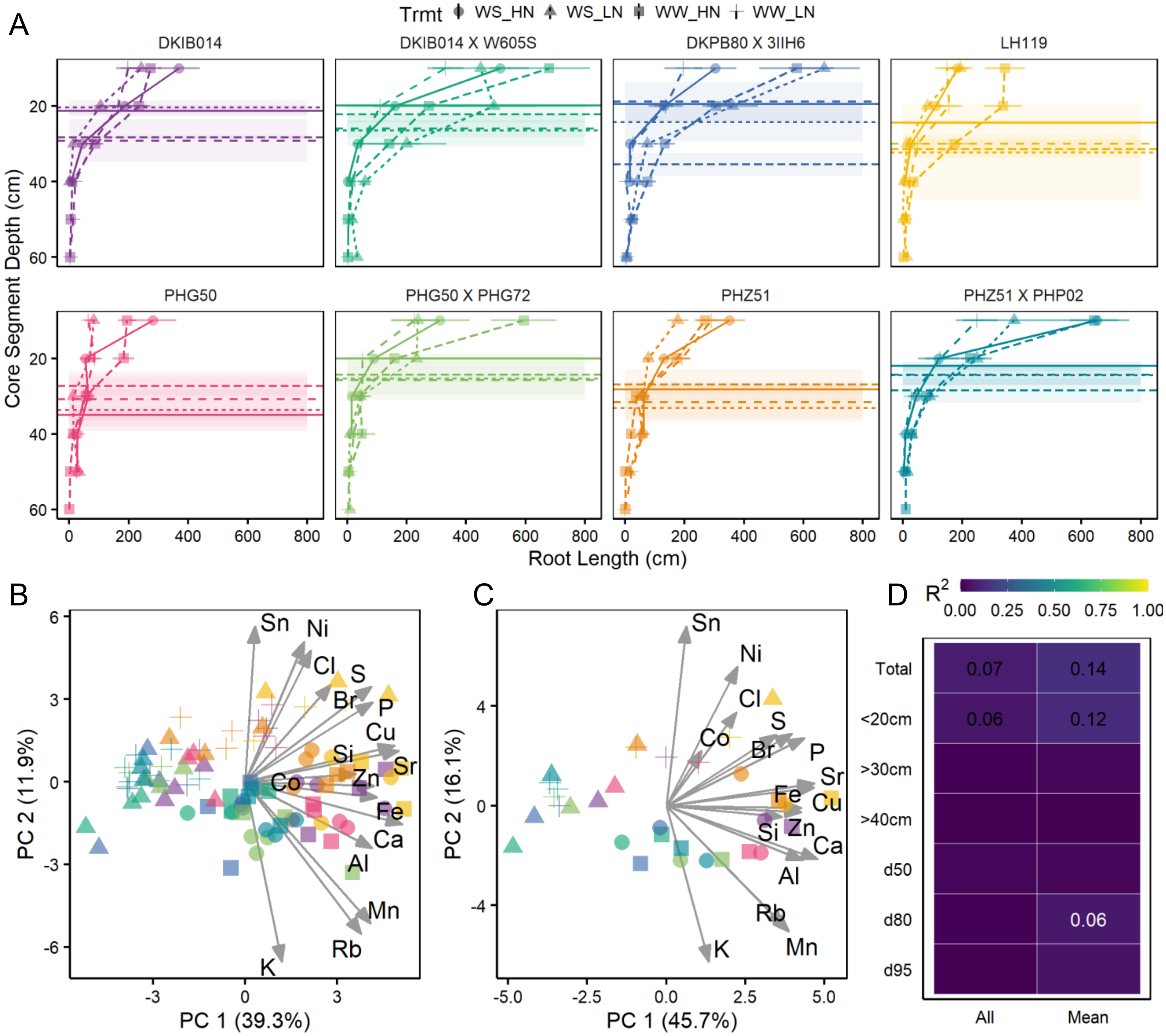


Supplemental Figure 7: HAN 2019 A) Root depth distributions of 8 genotypes grown in the field under well-watered (WW), water-stressed (WS), high-nitrogen (HN), and low-nitrogen (LN) treatment combinations. Circles represent WS_HN; triangles, WS_LN; squares, WW_HN; and crosses, WW_LN. Points represent the root length in each 10 cm layer and error bars represent the standard error of the mean. Dotted lines represent the D95 and shaded regions, the standard error. Color indicates weeks after planting. B) PCA biplots of all plants with triangles representing the WW treatment. Loadings on the PCs are indicated with labeled arrows. C) PCA biplot of mean elemental values of the treatments and genotypes for each treatment group. D) Correlations ($R^2$) between root traits (Y) and PC1 of either all individual plots or the mean of elemental values and root values. Correlation values are shown when they are significant at the $p \leq 0.05$ level in black and in white when $p \leq 0.1$.

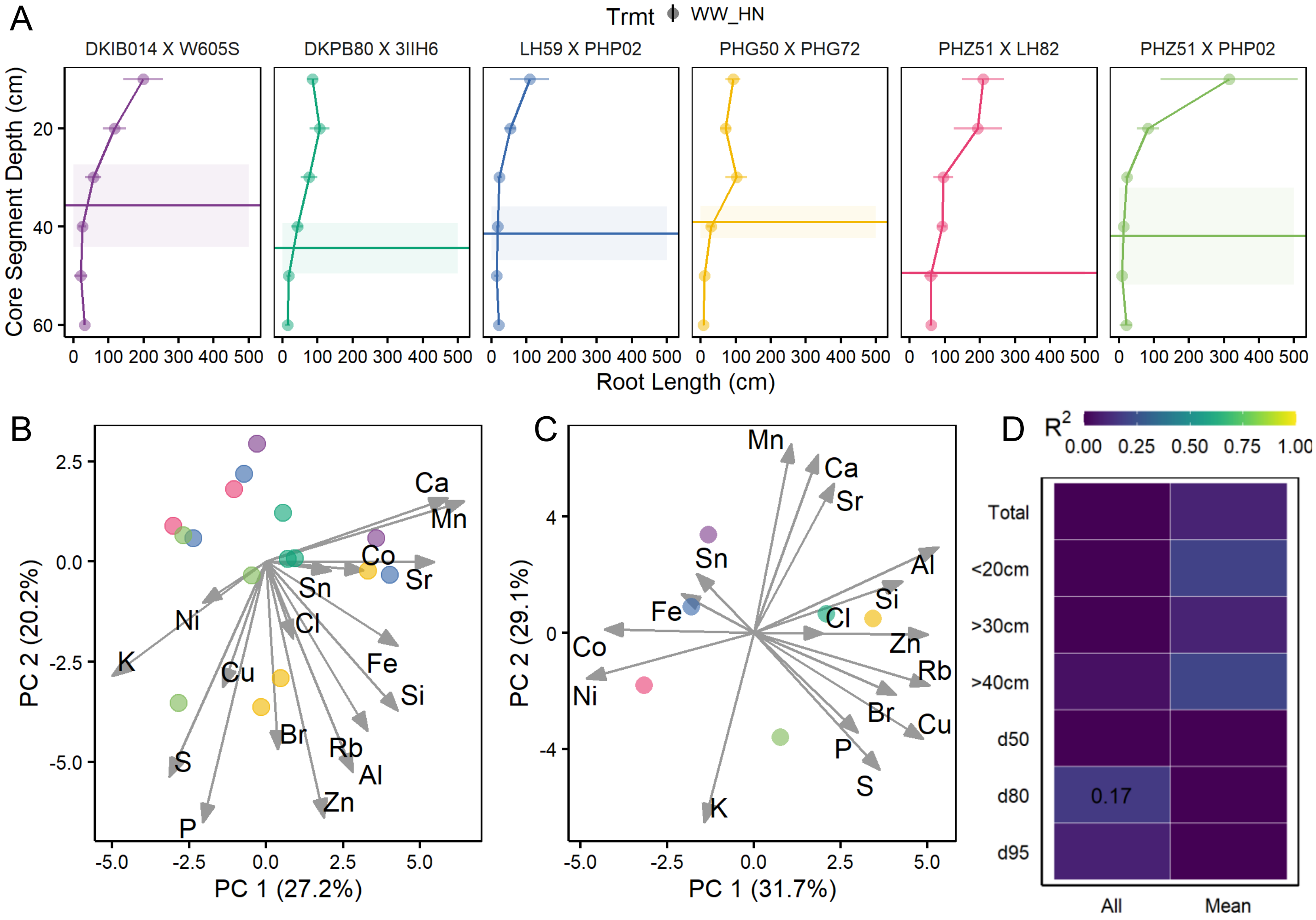


Supplemental Figure 8: ARL 2019 A) Root depth distributions of 6 genotypes grown in the field in Arlington, WI. Points represent the root length in each 10 cm layer and error bars represent the standard error of the mean. Dotted lines represent the D95 and shaded regions, the standard error. Color indicates weeks after planting. B) PCA biplots of all plants with triangles representing the WW treatment. Loadings on the PCs are indicated with labeled arrows. C) PCA biplot of mean elemental values of the treatments and genotypes for each treatment group. D) Correlations ($R^2$) between root traits (Y) and PC1 of either all individual plots or the mean of elemental values and root values. Correlation values are shown when they are significant at the $p \leq 0.05$ level.

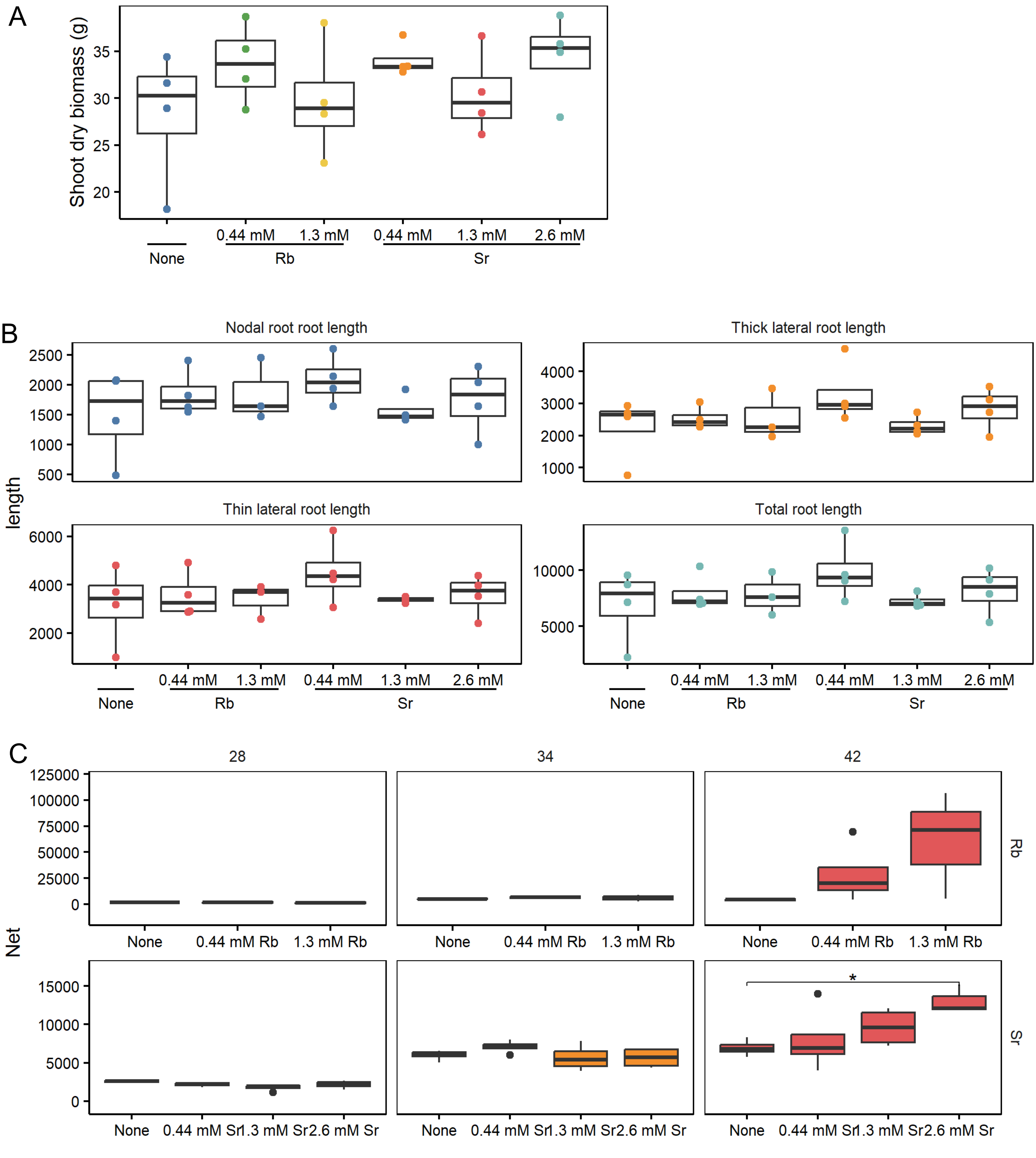


Supplemental Figure 9. A) Shoot dry biomass of greenhouse-grown maize plants after 48 days. Tracers for root depth (0.06M Sr – 0.17M Rb) did not impact biomass (ns, Dunn Test). B) Root length metrics of 48-day old greenhouse grown maize plants grown with tracers. There were no significant effects of tracers. C) XRF counts of the oldest, green leaf of plants with tracers applied. Only after 28 days were plants provided 0.34 M Sr showing significant accumulation, though we saw dose-dependent increasing trends ($p \leq 0.05$, t-test compared to control conditions).

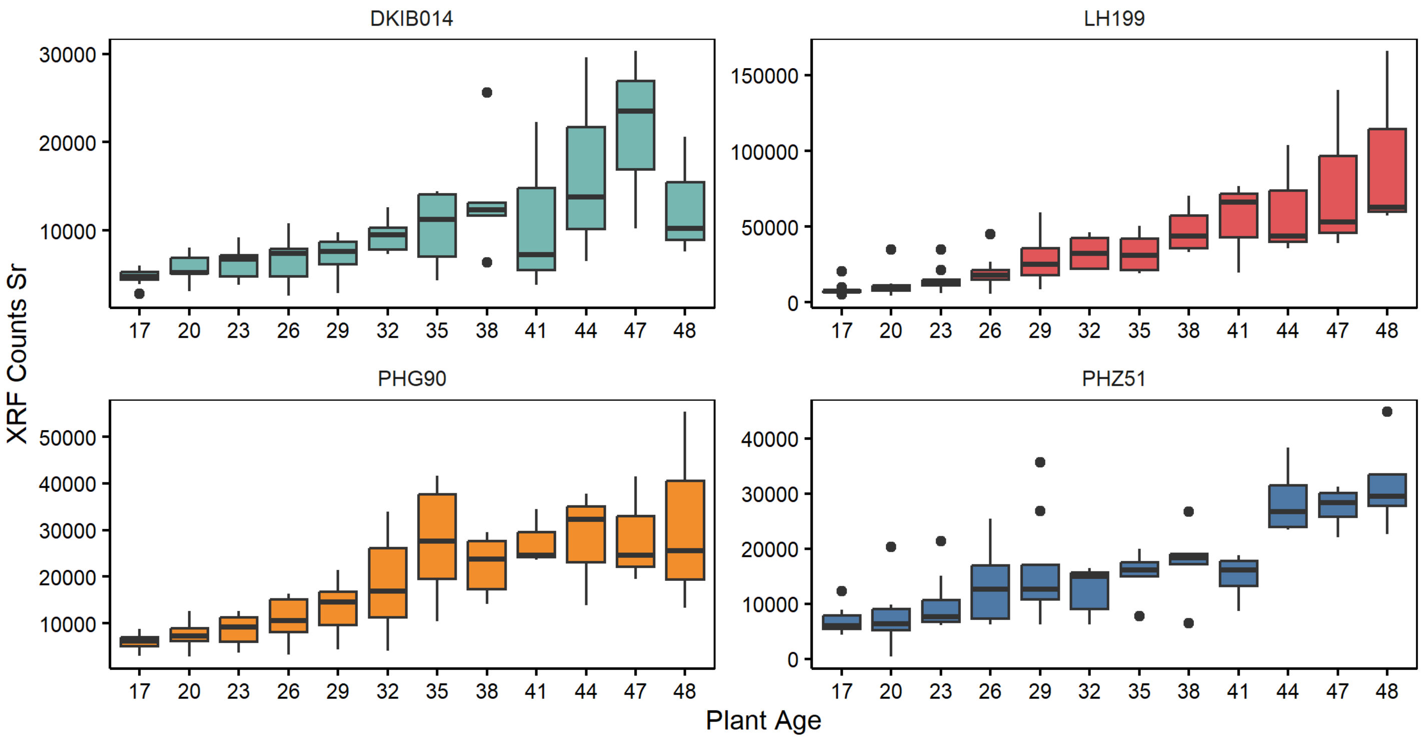


Supplemental Figure 10. Accumulation of Sr measured by XRF in the oldest green leaf over time in the greenhouse (2019 experiment). Four genotypes were measured (n = 9).
